# INS-17, an insulin-receptor ligand, confers specificity in adult IIS-regulated phenotypes in *C. elegans*

**DOI:** 10.1101/2025.11.20.689590

**Authors:** Emily J. Leptich, Priyadharshini Vijayakumar, Edward W. Pietryk, Meredith I. Williams, Rachel N. Arey

**Affiliations:** Department of Neuroscience, Baylor College of Medicine, Houston, Texas, United States of America; Center for Precision Environmental Health, Baylor College of Medicine, Houston, Texas, United States of America; Department of Molecular and Human Genetics, Baylor College of Medicine, Houston, Texas, United States of America; Department of Molecular and Cellular Biology, Baylor College of Medicine, Houston, Texas, United States of America

## Abstract

Insulin/IGF growth factor (IGF) signaling (IIS) is a pleiotropic signaling pathway that functions across tissues to coordinate phenotypic changes in response to nutrient status. Thus, the ubiquity of the IIS pathway hinders efforts to elucidate the mechanisms driving specific IIS-related phenotypes, Previous research in the nematode worm *C. elegans* has demonstrated that loss of function of the IGF transmembrane receptor (IR) ortholog, DAF-2, results in a doubled lifespan and enhanced learning and memory behaviors in young and aged animals. However, these findings are the result of reducing DAF-2 receptor function rather than modulating ligand-receptor interactions. In the current study, we aimed to dissect ligand-receptor interactions that may regulate associative behaviors apart from canonical IIS lifespan phenotypes in *C. elegans*. To this end, we performed targeted genetic screening of Insulin-like Peptides (ILPs) previously identified as DAF-2 antagonists to test their role in learning and memory phenotypes. We discovered that only a single uncharacterized ILP, INS-17, is required for learning and memory. We also demonstrate that INS-17 is sufficient to confer extended memory ability and can promote the maintenance of learning and memory with age. Additionally, we observe that *ins-17* regulates learning and memory ability independent of lifespan, uncoupling IIS-mutant phenotypes. We find that regulation of the *ins-17* genetic locus explains its unique requirement among ILP for learning and memory behaviors. Finally, we found that INS-17 may act to signal a state of nutrient deprivation that is required to properly process stimulus valence to promote advantageous behaviors. Our findings deepen the understanding of how IIS can regulate specific phenotypic outputs in response to changes in internal metabolic states.

## Introduction

The survival of most multicellular organisms depends on the coordination of complex physiological processes between multiple, functionally discrete tissues. Remarkably, each of these tissues carries out a diverse set of physiological functions using a highly overlapping set of cell regulatory mechanisms, including ubiquitous signaling pathways (e.g. MAPK/ERK, cAMP, and ubiquitin-proteasome pathway signaling(1–4)). Major components of these pathways are commonly pleiotropic, where a single gene can control multiple, distinct phenotypes depending on the context, such as life stage, cell type, and metabolic state. Across species, one prominent pleiotropic signaling pathway is the insulin/insulin-like growth factor (IGF) receptor signaling (IIS), which regulates a wealth of phenotypes across multiple tissues, including growth and development, metabolism, fertility, aging, stress response, and survival (5–15). Dissecting the specific signaling mechanisms by which IIS mediates such a wide range of phenotypic effects proves challenging, especially in mammalian systems. This is due to several factors, namely high costs to generate tools to examine pathways in a tissue-specific and temporally controlled manner; compensation between receptors, as a single tissue can regulate phenotypes organism-wide; and the time and resource burden to study certain IIS-regulated phenotypes, such as aging. However, invertebrate systems can help overcome these challenges, as IIS is highly conserved across species (9,10,14,16,17), and simple models allow for rapid, low-cost study of genes with remarkable temporal and tissue-specific precision (18–20).

The nematode *C. elegans* has been invaluable in elucidating the pleiotropy underlying IIS phenotypes. In fact, foundational experiments in the worm were the first to link insulin signaling and aging, where loss-of-function receptor mutants of the IR homolog, *daf-2*, have a doubled lifespan of around 40 days compared to the 20-day lifespan of wild-type animals (21,22). Since this initial work, studies in IIS mutants revealed the pleiotropy of the DAF-2 receptor and its conserved downstream signaling components (*age-1*/PI3K and transcription factors *hsf-*1/HSF1, *skn-1*/NRF2, and *daf-16*/forkhead family (FOXO3)) in this relatively simple model (21–28), including that *daf-2* mutants have altered development, metabolism, fertility, motility, cognitive abilities, and more (12,26,29–39). Collectively, research indicates that DAF*-2/*IR functions across tissues and is integral for sensing internal state along with environmental cues allowing for the worm to adapt appropriately to improve survival.

Despite decades of study of IIS in the worm, how the DAF-2 receptor coordinates a variety of phenotypic responses across tissues remains unclear. This is due in part to the ubiquity of the IIS pathway across tissues, where DAF-2/IR activity in some tissues can mask DAF-2/IR activity in other tissues. For example, it is well established that *daf-2* mutants have improved motility with age (30,40,41), but recent work using degron-based approaches found that reduction of *daf-2* function in neurons and muscles has opposing effects on maintenance of motility in aged animals (42). This work sheds light on the remaining gap in our knowledge regarding how signaling through a single ubiquitous receptor can impact numerous phenotypes that are often reflective of the function of a relatively small number of cells. It is also unclear whether these distinct phenotypes can be uncoupled, and if so by what mechanisms.

Though the DAF-2 receptor and its downstream effectors have been well-studied, less is known about the upstream regulators of DAF-2/IR that drive specific phenotypic outputs. The *C. elegans* genome encodes 40 insulin-like peptides (ILPs) predicted to act as DAF-2 receptor agonists or antagonists, sometimes in a context-dependent fashion (43–47). However, no single manipulation of an ILP fully reproduces reduced *daf-2* function phenotypes. ILPs also exhibit diverse and often non-overlapping expression patterns and are regulated by distinct physiological contexts (48). Thus, we hypothesize that pleiotropic phenotypes exhibited by IIS mutants may also reflect distinct ligand-receptor interactions.

In the present study, we investigated if specific ILPs regulate *daf-2* mutant learning and memory phenotypes. We discovered that a single ILP, the DAF-2/IR antagonist INS-17, is required for learning and memory ability and is sufficient to promote memory. We also found that INS-17 is specifically required for IIS regulation of behavior and does not appear to be involved in other adult DAF-2 receptor-mediated phenotypes, including lifespan (47,49). We find that the regulation of INS-17 is the source of this phenotypic specificity, and that INS-17 acts as a nutrient-deprivation signal that regulates plastic behavior in the context of changing physiological states.

## Results

### INS-17 is specifically required for learning and memory ability

Decreased DAF-2 receptor function results in a 3-fold extension in short- and intermediate associative memory (S/ITAM), as measured by positive olfactory association assays (31). In this paradigm, animals form a positive association with the neutral odor butanone (10% in EtOH) when paired with food, which is measured by a training-dependent increase in butanone chemotaxis (50). A single food-butanone (1 CS-US) pairing usually results in a molecularly conserved memory that lasts approximately 2 hours; however, young adult *daf-2* mutants maintain a food-butanone association for up to 6 hours (Fig. 1A) (31). Percent maximum (% Max) performance indices are shown for clarity (see methods for calculations), as *daf-2* animals have high naïve chemotaxis to 10% butanone in multiple studies (31,34,36,37) (Fig. S1A) that contribute to perceived learning impairments when graphed with raw performance indices (provided in Fig. S1B). Given that partial loss-of-function *daf-2* animals display this significant extension in S/ITAM performance, we hypothesized that insulin receptor antagonist(s), which are predicted to reduce signaling through DAF-2/IR, may promote learning and memory behavior (Fig. 1B). To test this hypothesis, we examined the learning and memory ability of publicly available insulin receptor antagonist mutants using this STAM paradigm. We tested learning and intermediate-term associative memory (ITAM, 60 minutes post-training), as behavioral deficits are typically more prominent at this timepoint compared to STAM (30 minutes post-training). Mutants tested included predicted strong (*ins −17, −37, and −39)* and weak (*ins-15, −21,* and *-22)* antagonists (47,51,52). However, we recently determined that mutants for *ins-22* have an abnormally high preference for 10% butanone (Fig. S1I) (53). Thus, we excluded *ins-22* from our targeted behavioral screen, as this high naïve chemotaxis would make it difficult to measure training-dependent increases in butanone preference. Most antagonist mutants behaved in a manner that was indistinguishable from wild-type animals, including baseline naïve chemotaxis (Fig. 1D-E, S1E-H); while mutant animals for *ins-39* had significantly higher intermediate-term associative memory (ITAM) performance compared to wild-type animals, we were screening for behavioral deficits and focused on candidates that had impaired behavioral performance. Interestingly, only loss-of-function mutants of *ins-17* exhibited behavioral deficits: these include significant deficits in learning, STAM and ITAM performance (Fig 1C). These deficits were not accompanied by impaired sensory function, as measured by naïve chemotaxis assays to the highly attractive concentration (0.1%) of butanone (Fig. S1C-D), suggesting that they are specific to learning and memory. INS-17 is a relatively uncharacterized ILP; aside from its prediction as a strong antagonist, previous research has established *ins-17’*s role in regulating dauer formation in early development (49). To our knowledge, INS-17 is uncharacterized regarding complex behavior, and these results (Fig. 1C-E) suggest that it is a novel learning and memory-regulating ILP.

**Fig. 1.**
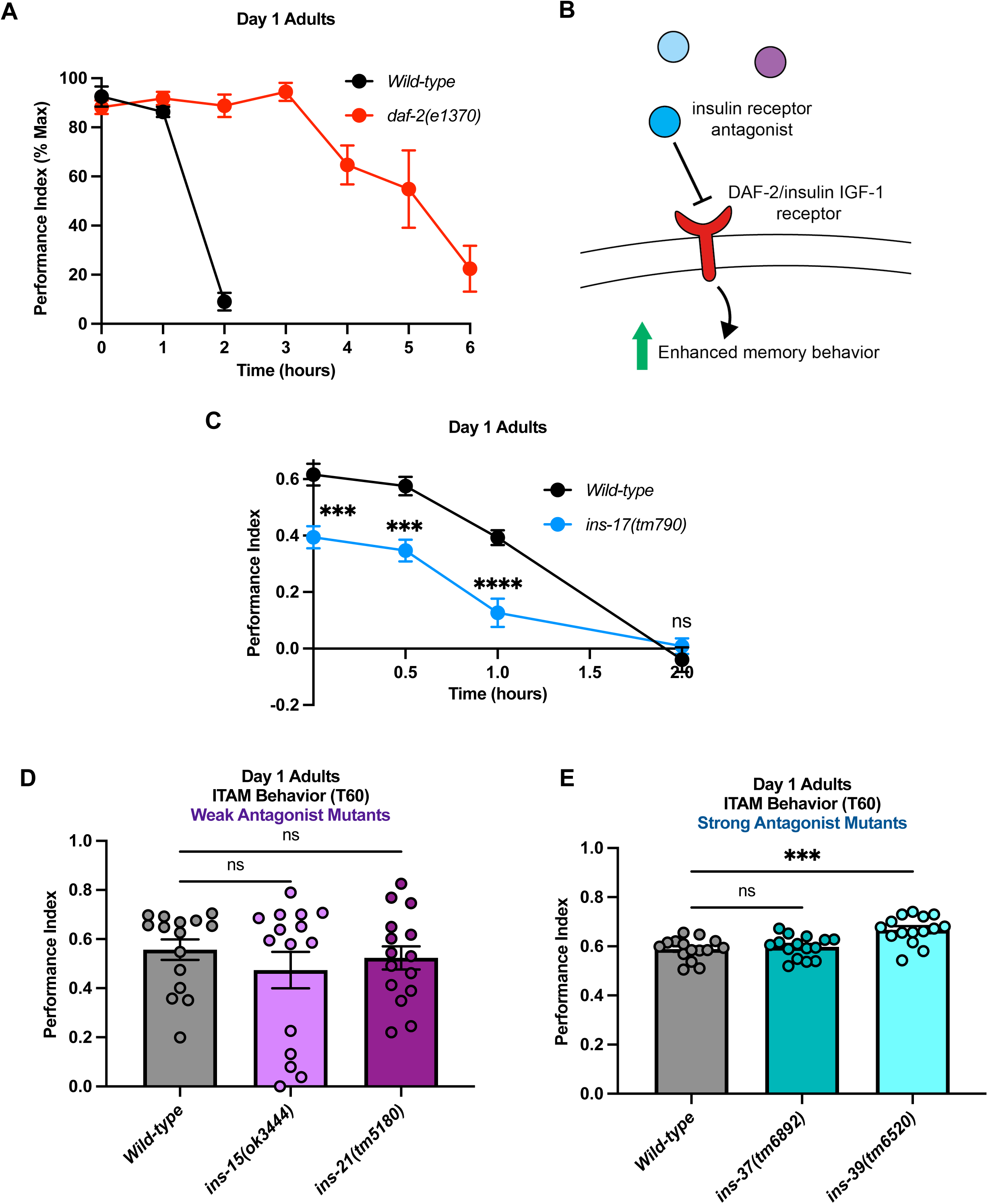
INS-17 is a novel and specific memory-regulating insulin receptor antagonist. **(A)** Partial loss-of-function *daf-2(e1370)* mutants have extended STAM that lasts 6 hours compared to 2 hours for wild-type animals after 1 CS-US food-butanone pairing. % Max Performance reported as *daf-2(e1370)* animals have a higher baseline naïve attraction to neutral (10%) butanone concentration (see **Fig. S1A, B**). Mean ± SEM. n = 15 per genotype. **(B)** Diagram depicting the hypothesis that insulin receptor antagonists may inhibit DAF-2 receptor function to promote learning and memory. **(C)** *ins-17* deletion mutants have impaired learning, STAM, and ITAM compared to wild-type worms. Mean ± SEM. n = 15 per genotype. ***p < 0.001, ****p < 0.001; ns, not significant (p > 0.05). **(D)** Weak insulin antagonist mutants for *ins-15* and *ins-21* and **(E)** strong insulin antagonist mutants for *ins-37* and *ins-39* display no memory impairments; *ins-39* mutants have better ITAM performance compared to wild-type. Mean ± SEM. n = 15 per genotype. ***p < 0.001.

### INS-17 signals through DAF-2 and promotes memory

We next wanted to confirm that INS-17 signaling was indeed mediated by the DAF-2 receptor and tested the learning and memory behavior of *daf-2(e1370);ins-17(tm790)* double mutants. We examined double mutant memory behavior at 3 hours after 1 CS-US pairing, a timepoint where *daf-2* mutants exhibit extended memory (31). *daf-2(e1370);ins-17(tm790)* worms demonstrated extended 3-hour memory indistinguishable from *daf-2* animals (Fig. 2A, S2A-C). Thus, we infer from these results that DAF-2/IR is epistatic to INS-17 in the regulation of learning and memory.

**Fig. 2.**
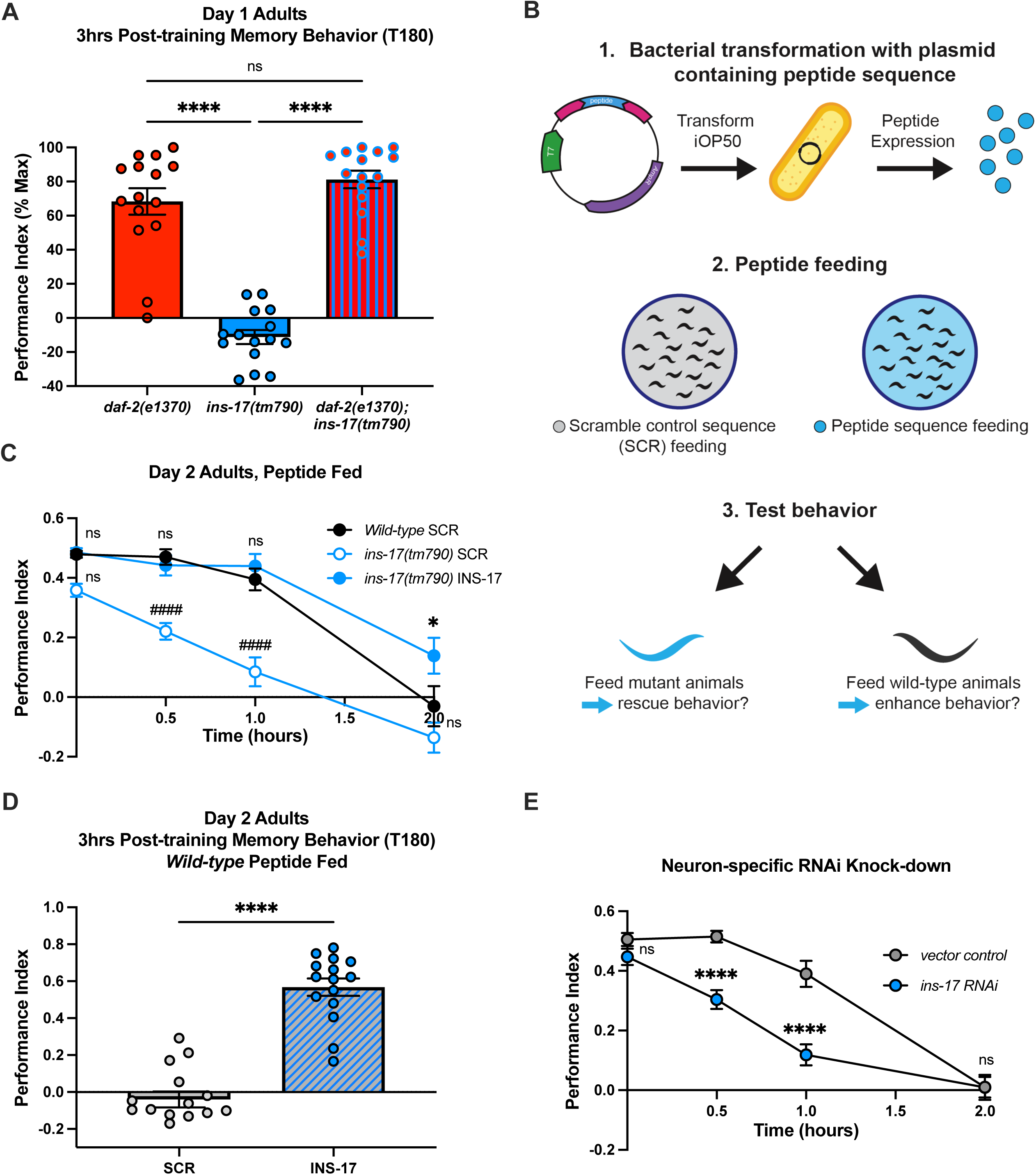
INS-17 and DAF-2 are epistatic and promote memory. **(A)** *daf-2* and *ins-17* share an epistatic relationship. 3hrs post-training (1 CS-US pairing), *daf-2(e1370)* mutants and *daf-2(e1370)*;*ins-17(tm790)* double mutants display memory behavior while *ins-17(tm790)* animals display no memory behavior. Mean ± SEM. n = 15 per genotype. ****p < 0.0001. **(B)** Diagram of microbial feeding paradigm. iOP50 bacteria were transformed with a plasmid containing either the peptide sequence of interest or a scramble control sequence and then fed to worms before testing behavioral effects. **(C)** INS-17 peptide feeding rescues *ins-17(tm790)* learning, STAM and ITAM behaviors, while SCR fed mutants display significant S/ITAM impairments compared to SCR fed wild-type animals. Mean ± SEM. n = 15 per feeding treatment. # is significance relative to SCR fed wild-type animal performance. ###p < 0.001, ####p < 0.0001; ns, not significant (p > 0.05). **(D)** Wild-type animals fed INS-17 peptides display 3-hour memory behavior after 1 CS-US pairing, similar to *daf-2(e1370)* extended STAM performance. Mean ± SEM. n = 15 per peptide treatment. ****p < 0.0001. **(E)** Adult-only, neuron-specific RNAi knock-down of *ins-17* leads to learning, STAM, and ITAM impairments compared to vector RNAi treated animals. Mean ± SEM. n = 15 per RNAi treatment. ****p < 0.0001.

Next, we wanted to explore how manipulating levels of INS-17 may influence behavioral outputs. Our behavior paradigm tests an adult-specific behavior (34), and *ins-17* regulates dauer in development (49), so we sought to avoid confounding our results with developmental effects of traditional transgenic approaches. Recently, we developed a novel microbial feeding paradigm to manipulate neuropeptide signaling specifically in adult animals (53). In these experiments, *C. elegans* are fed *E. coli* bacteria expressing a plasmid containing the sequence encoding a neuropeptide of interest flanked by processing enzyme cut sites (Fig. 2B). Like feeding-based RNAi approaches (54), this method offers precise temporal control of increasing neuropeptide levels via adult-only manipulation to avoid potential effects on early development. We have demonstrated that this feeding method could work in an ‘overexpression’ manner when administered to wild-type animals, altering sensory behavior (53). Here, we wanted to extend this approach to assess if administering additional INS-17 to adult animals could affect associative behaviors. We hypothesized that increasing the levels of a memory-regulating *daf-2* antagonist would result in enhanced memory ability. Briefly, we fed L4 animals *E. coli* expressing either INS-17 or a scrambled control peptide (SCR) and examined their behavior at Day 2 of adulthood.

To first establish the ability of this method to effectively manipulate INS-17 signaling, we tested if INS-17 peptide feeding was sufficient to rescue learning and memory behavior of *ins-17(tm790)* animals. SCR fed mutants still exhibited behavioral deficits, while adult-only feeding of INS-17 to *ins-17* deficient mutants was sufficient to rescue their behavior, similar to wild-type animals fed SCR peptide (Fig. 2C). Notably, we determined these results were not the result of sensory deficits in these animals (Fig. S2D-E). We next explored if INS-17 peptide feeding was sufficient to boost wild-type animal memory behavior past the 2-hour timepoint after 1 CS-US pairing. Remarkably, we found that wild-type animals fed INS-17 during adulthood displayed 3-hour memory post-training, while SCR fed animals demonstrated no memory phenotype, as expected (Fig. 2D). We confirmed that peptide feeding to both wild-type and mutant animals did not alter baseline chemosensation, indicating the effect is memory-specific (Fig. S2D-F). These results suggest that increasing INS-17 is sufficient to boost cognitive performance and phenocopy *daf-2* animals’ 3-hour memory behavior.

Finally, we worked to identify the tissue where INS-17 is required to regulate memory. Since INS-17 is highly expressed in the nervous-system, we hypothesized that neurons may use INS-17 as a memory signal. Using a neuron-specific RNAi-sensitive line(55), we performed an adult-only *ins-17* knockdown and observed significant S/IATM behavioral deficits in *ins-17 RNAi* knock-down animals (Fig. 2E)—similar to what we observed in deletion mutants (Fig. 1B)—with no detectable effects on sensory ability (Fig. S2H-I). This demonstrates that INS-17 is a memory-promoting DAF-2/IR antagonist, likely released from neurons.

### INS-17 regulates the cognitive aging phenotypes mediated by the DAF-2 receptor independent of lifespan

In addition to exhibiting extended STAM at young adult stages, *daf-2* mutants also maintain the ability to learn and remember after 1 CS-US pairing with age (31). Wild-type animals display age-related cognitive deficits as early as Days 3 and 5 of adulthood (Fig 3A-B, S3A-C), while *daf-2* mutants maintain performance akin to Day 1 of adulthood at these timepoints. Therefore, we wanted to ask if INS-17 could also be involved in these age-related phenotypes associated with altered *daf-2* function. As *ins-17* mutants already display severe behavioral defects at Day 1 of adulthood, we examined the effect of increased INS-17 on age-related behavioral impairments in wild-type animals, again using a peptide feeding-based approach (53).

**Fig. 3.**
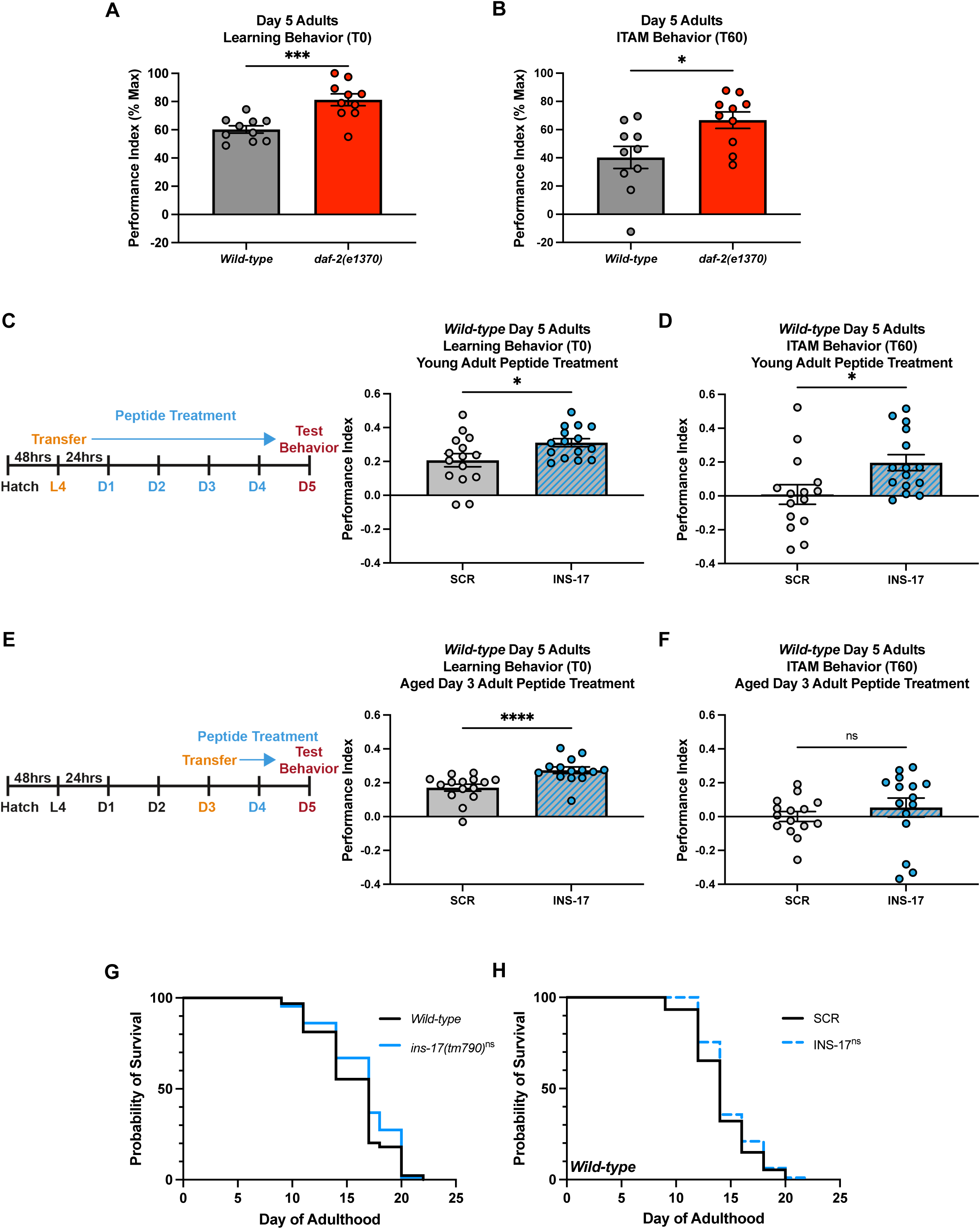
INS-17 regulates cognitive aging phenotypes observed in *daf-2* mutants. **(A)** Day 5 adult *daf-2(e1370)* mutants display better learning and **(B)** memory behavior compared to wild-type worms. Mean ± SEM. n = 10 per genotype. *p < 0.05, *** p < 0.001. **(C)** Timeline of young adult peptide treatment. Wild-type L4 animals are fed SCR or INS-17 iOP50 bacteria until Day 5 of adulthood when S/ITAM behavior is tested. INS-17 peptide treated worms have increased learning and **(D)** ITAM performance compared to SCR peptide treated animals. Mean ± SEM. n = 15 per peptide treatment. *p < 0.05. **(E)** Timeline of aged adult peptide treatment. Wild-type Day 3 adult animals are fed SCR or INS-17 iOP50 bacteria until Day 5 of adulthood when S/ITAM behavior is tested. INS-17 peptide treated worms have increased learning performance, but not **(F)** ITAM performance, compared to SCR peptide treated animals. Mean ± SEM. n = 15 per peptide treatment. *p < 0.05, ****p < 0.0001. **(G)** Probability of survival analysis demonstrates there is no significant difference in lifespan comparing wild-type animals and *ins-17(tm790)* animals. One replicate depicted is representative of three replicates (see **Fig. S3F, S3G**). Mean/median ± SEM. n ≥ 100 per replicate. ns, not significant (p > 0.05). **(H)** Probability of survival analysis for wild-type animals treated with SCR peptide compared to wild-type animals fed INS-17 peptide shows no significant difference in average lifespan. One replicate depicted is representative of three replicates (see **Fig. S3H, S3I**). Mean/median ± SEM. n ≥ 100 per replicate. ns, not significant (p > 0.05).

We first assessed the behavior of animals supplemented continuously with INS-17 from the L4 stage until Day 5 of adulthood (Fig. 3C), when worms are considered cognitively aged (31). INS-17 fed animals demonstrated significantly better learning and memory performance compared to wild-type SCR fed controls (Fig. 3C-D, S3D). Next, we wondered if manipulating the DAF-2/IR pathway via INS-17 later in life could influence cognitive aging, as previous work has shown that degradation of DAF-2/IR via an auxin inducible degron system in older animals results in improved STAM (37). We subjected Day 3 adult wild-type animals (which already display age-related cognitive deficits compared to Day 1 Adults (Fig. 3A)) to SCR or INS-17 peptide feeding and tested their learning and memory abilities at Day 5 of adulthood (Fig. 3E). Notably, we found that Day 5 INS-17 fed animals had better learning and STAM behaviors compared to SCR fed controls (Fig. 3E-F, S3E-F). These findings suggest that INS-17 is a DAF-2/IR ligand that could partially contribute to the enhanced cognitive ability with age observed when DAF-2/IR activity levels are reduced.

Since *daf-2* animals have both a doubled lifespan and enhanced memory phenotypes in young and aged animals (21,31), we wanted to determine if manipulating INS-17, as a strong DAF-2/IR antagonist, would alter lifespan in addition to associative behaviors. As previously reported in the literature (49), we confirmed that *ins-17(tm790)* animals have no significant difference in lifespan to wild-type animals (Fig. 3G, S3F-G). We also tested the lifespans of wild-type animals fed SCR compared to those fed INS-17 in adulthood, finding no significant differences between groups (Fig. 3H, S3H-I). Thus, our findings demonstrate that either loss of *ins-17* or increased levels of INS-17 has no effect on adult lifespan, highlighting the specific phenotypic regulation by individual ligands underlying the pleiotropy of the DAF-2 receptor.

### INS-17 engages multiple pathways downstream of DAF-2 to regulate learning and memory

We next sought to identify the pathways downstream of DAF-2/IR that could explain the phenotypic specificity of INS-17. The canonical downstream target of DAF-2/IR signaling is the FOXO transcription factor homolog DAF-16(27,28), which is required for the doubled lifespan extension of *daf-2* loss-of-function worms (26,28,56). Additionally, *daf-16* mutants exhibit severely impaired learning and memory phenotypes (Fig. 4A, S4A (31). Since *ins-17* is required for positive associative memory behavior and there are no adult lifespan phenotypes associated with *ins-17* (Fig 3G-H), we hypothesized that *ins-17* activates a pathway that acts in parallel to *daf-16*. To test this, we examined if INS-17 feeding could improve the memory impairments of *daf-16* worms. We found that adult-only INS-17 feeding rescued the learning and memory of *daf-16(mu86)* animals to SCR fed wild-type performance levels, while SCR fed *daf-16(mu86)* worms still displayed behavioral deficits (Fig. 4B, S4B-C). These results support that INS-17 could act independently of DAF-16 to regulate associative behaviors.

**Fig. 4.**
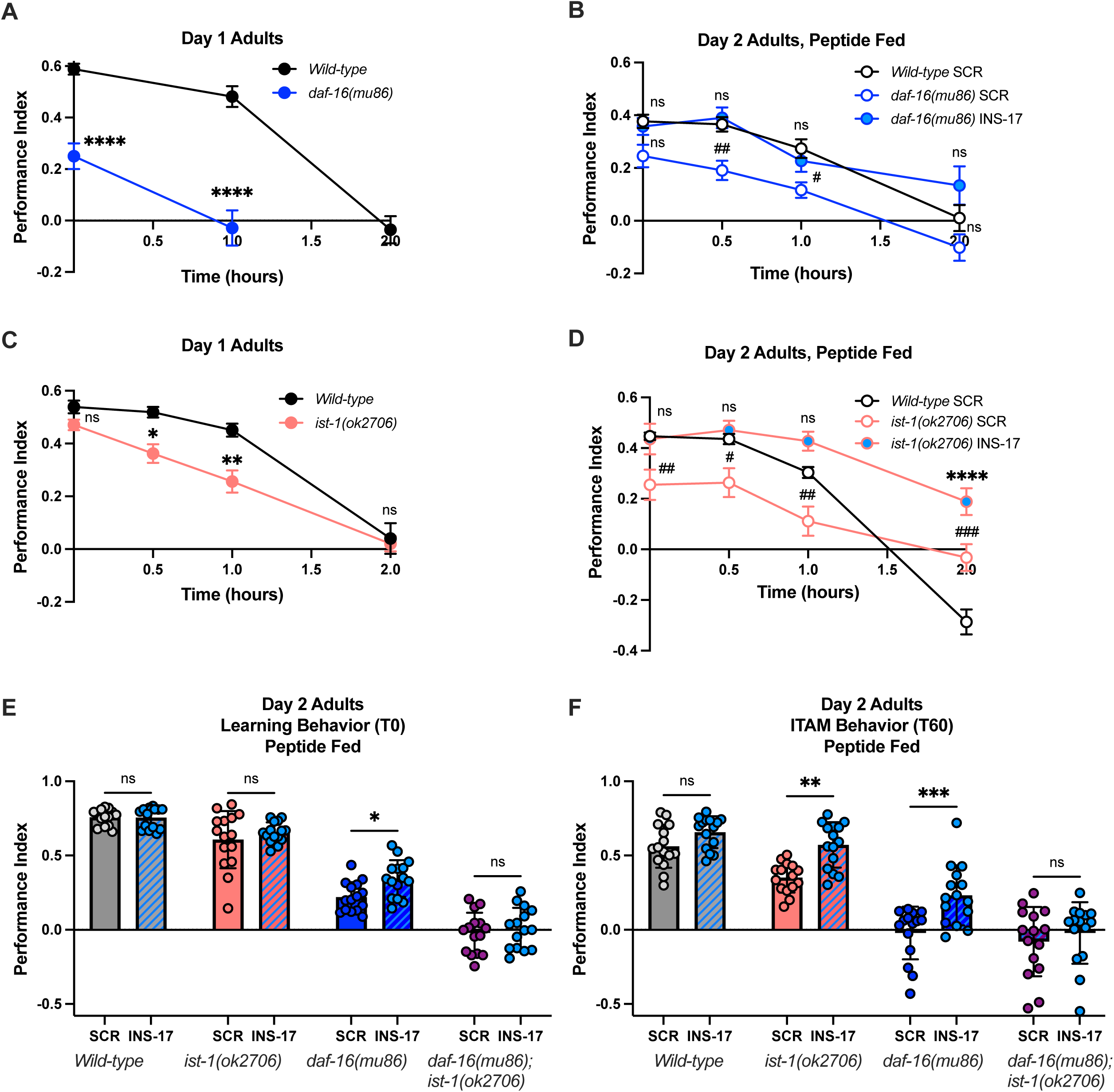
INS-17 likely engages parallel pathways of DAF-2 receptor signaling. **(A)** *daf-16(mu86)* deletion mutants have impaired learning and memory behavior compared to wild-type animals. Mean ± SEM. n = 15 per genotype. ****p < 0.0001. **(B)** INS-17 peptide treatment rescues *daf-16(mu86)* S/ITAM impairments. Mean ± SEM. n = 15 per genotype and peptide treatment. Wild-type animals fed INS-17 performance indices are in **Fig. S4C**. # is significance relative to wild-type. #p < 0.05, ##p < 0.01; ns, not significant (p > 0.05). **(C)** *ist-1(ok2706)* deletion mutants display a slight but not significant learning deficit and impaired S/ITAM behaviors compared to wild-type worms. Mean ± SEM. n = 15 per genotype. *p < 0.05, **p < 0.01; ns, not significant (p > 0.05). **(D)** INS-17 peptide treatment rescues *ist-1(ok2706)* learning and memory impairments. Mean ± SEM. n = 15 per genotype and peptide treatment. # is significance relative to wild-type. #p < 0.05, ##p < 0.01, ###p < 0.001, ****p< 0.0001; ns, not significant (p > 0.05). **(E)** *daf-16(mu86)*;*ist-1(ok2706)* animals display significant learning and **(F)** memory impairments that are not rescued by INS-17 peptide feeding, while *daf-16(mu86)* and *ist-1(ok2706)* deficits are rescued by INS-17 peptide administration. Mean ± SEM. n = 15 per genotype and peptide treatment *p < 0.05, **p < 0.01, ***p < 0.001; ns, not significant (p > 0.05).

Interestingly, recent research has uncovered evidence of pathways that can regulate cognitive abilities downstream of DAF-2/IR in parallel to DAF-16. This includes the discovery of a novel role for the insulin receptor substrate *ist-1*, which has been shown to function partially in parallel to DAF-16 to regulate learning ability in an aversive olfactory training paradigm (57). Thus, we aimed to determine if *ins-17* is functioning in the same pathway as *ist-1*. First, we examined if *ist-1* is required in our positive associative memory paradigm. We found that *ist-1(ok2706)* animals have impaired positive associative STAM behavior compared to wild-type animals (Fig. 4C). Like *daf-16* animals, we performed adult-only INS-17 feeding with *ist-1(ok2706)* worms and found that INS-17 peptide feeding was sufficient to boost STAM performance similar to wild-type animals fed SCR, while *ist-1* animals fed SCR still displayed deficits (Fig. 4D). Furthermore, we confirmed *ist-1* deficits were not the result of altered baseline chemosensation of butanone nor peptide feeding (Fig S4D-G).

While IST-1 and DAF-16 can function in parallel, it is possible that these two pathways can compensate for one another. Indeed, Cheng et al., (2022) identified that while *ist-1* could regulate learning independently of *daf-16*, *daf-16* was still required for this behavior(57). We hypothesized that INS-17 could similarly engage both *daf-16* and *ist-1* to promote memory ability, and that partial regulation of either arm of the DAF-2 receptor signaling pathway could act as a compensatory mechanism in the mutant backgrounds when fed INS-17. Therefore, we examined if the beneficial effects of INS-17 administration were abolished when both genes were mutated. We generated *daf-16(mu86);ist-1(ok2706)* double mutants and observed that they had significant learning and memory deficits (Fig. 4E-F) compared to either single mutant alone. Because of the striking nature of these deficits, we confirmed that double mutant animals still maintained chemosensory ability and determined that they exhibited normal preference for chemoattractants (Fig. SH-I). We found that INS-17 peptide administration failed to rescue these severe behavioral defects, in contrast to the improvement observed in single mutants (Fig. 4E-F, S4J-K). These results suggest that INS-17 feeding can activate DAF-16 and IST-1, partially in parallel, to regulate associative behaviors.

### Regulation of the *ins-17* genetic locus is required for proper regulation of behavior

While we had identified how INS-17 might regulate distinct DAF-2/IR mediated phenotypes, we also sought to examine what made INS-17 unique relative to other antagonists in regard to regulating behavior. Interestingly, though the *C. elegans* genome encodes 40 ILPs, there are only three structurally similar antagonists known as γ-insulins; these genes are encoded across multiple genomic regions (ch. II and III) and thus likely under the control of distinct regulatory elements (43,47,57). We therefore hypothesized that phenotypic specificity was not necessarily due to insulin class or structure, but instead via *ins-17*’s regulation by its surrounding genomic elements, which likely respond to different signaling pathways and environmental cues. To test this hypothesis, we explored if replacing *ins-17* with another strong DAF-2/IR antagonist was sufficient to maintain normal positive associative memory ability in an *ins-17* mutant background.

We selected *ins-37* to swap in for *ins-17* due to a key overlap in structure, as both ILPs are predicted strong DAF-2/IR antagonists and are γ-insulins, which are defined by three di-sulfide bonds (47). Moreover, *ins-37* mutants had no observable learning and memory impairments (Fig. 1D, S1H) and *ins-37* is lowly expressed in neurons (58), making it an ideal strong antagonist to test our hypothesis. Thus, we generated an *ins-17* CRISPR knockout in which we placed the *ins-37* coding sequence at the *ins-17* locus (Fig. 5A). Subsequently, we examined if *ins-37* expressed at endogenous *ins-17* levels is sufficient for normal learning and memory behavior. First, we confirmed that the newly generated *ins-17* knockout line exhibited learning and memory impairments similar to the predicted loss-of-function mutants (Fig 1C, 5B, S5A). Interestingly, we found that replacing *ins-17* with *ins-37* could rescue the defective learning and memory of *ins-17* KO animals, functionally replacing endogenous *ins-17* (Fig. 5B, S5A). These results indicate that ILP structure is not as critical as its regulation of expression. These results suggest that the genomic context in which *ins-17* is regulated underlies its importance in promoting specific behavioral phenotypes.

**Fig. 5.**
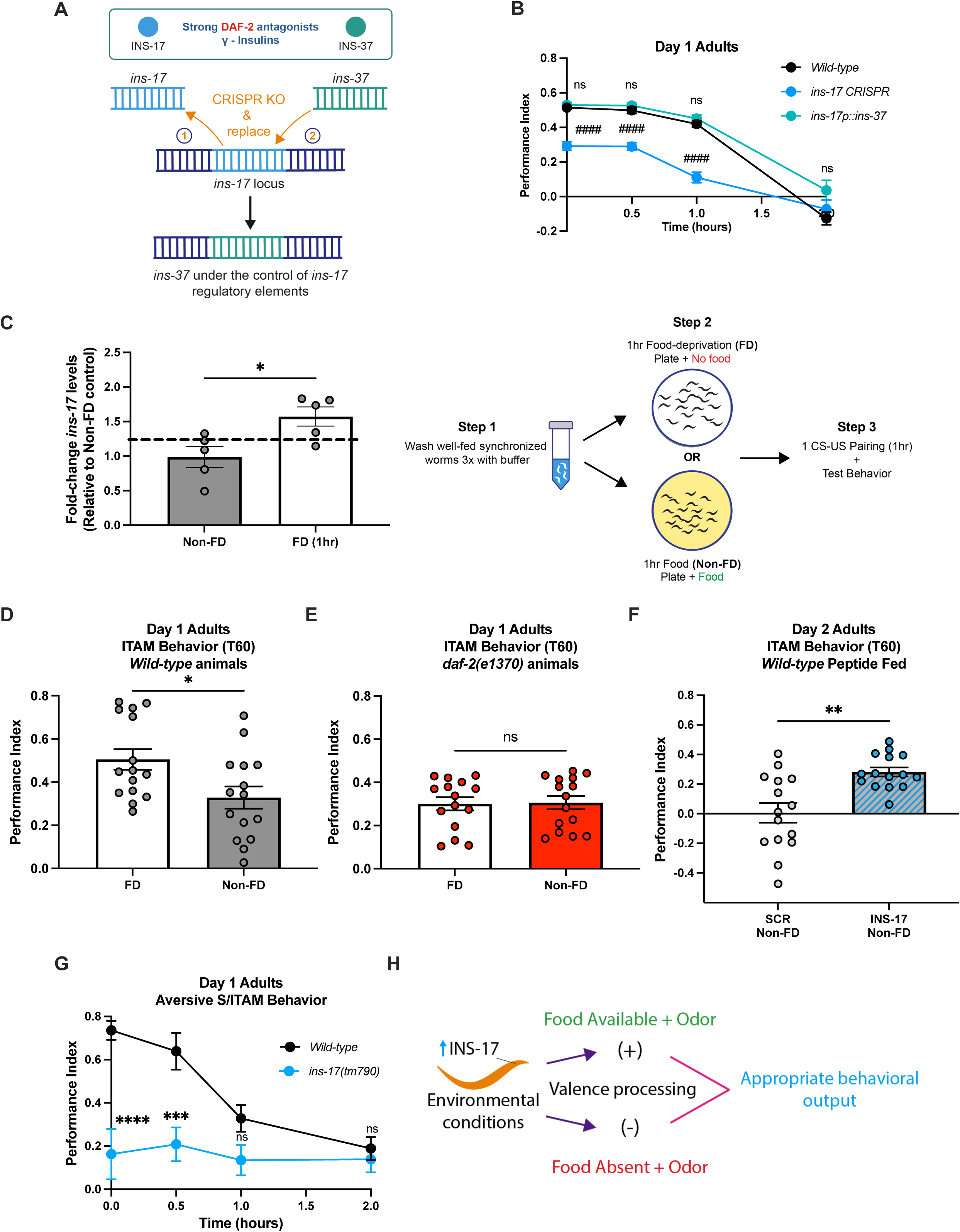
INS-17 is regulated by nutrient deprivation to promote advantageous behaviors. **(A)** Diagram showing *ins-17* replacement experiment design. *ins-17* and *ins-37* encode γ -ILPs. *ins-17* was removed with CRISPR and replaced with *ins-37* at the *ins-17* locus so *ins-37* would be expressed at endogenous *ins-17* levels. **(B)** Replacing *ins-17* with *ins-37* has no significant effect on behavior compared to wild-type animals, while *ins-17* CRISPR KO (*ins-17(knu1320)*) animals have impaired learning and S/ITAM. Mean ± SEM. n = 15 per genotype. # is significance relative to wild-type. ####p < 0.0001; ns, not significant (p > 0.05). **(C)** qRT-PCR of *ins-17* mRNA levels in Day 1 adult wild-type animals show that one hour of food-deprivation (FD) results in increased *ins-17* transcript levels (left). For levels measured after two hours of food-deprivation, please see **Fig. S5B.** Schematic depicting food-deprivation experiment workflow (right): well-fed adult worms are washed three times in M9 buffer solution to remove bacteria and placed on 100mm agar plates either seeded with a bacterial lawn (non-FD) or no lawn (FD) prior to training and behavior testing. Mean ± SEM. n = 5 per genotype. *p < 0.05. **(D)** Wild-type animals display more robust ITAM behavior if a FD step is administered prior to 1 CS-US pairing compared to training alone (non-FD conditions). Mean ± SEM. n = 15 per condition. *p < 0.05. **(E)** Food-deprivation prior to training (1 CS-US pairing) does not enhance *daf-2(e1370)* ITAM performance. Mean ± SEM. n = 15 per condition. ns, not significant (p > 0.05). **(F)** INS-17 peptide treatment in wild-type animals is sufficient to promote robust ITAM performance in non-FD conditions prior to training compared to non-FD SCR fed animals. See **Fig. S5I** for food-deprived SCR and INS-17 peptide fed comparisons. Mean ± SEM. n = 15 per peptide treatment. **p < 0.01. **(G)** *ins-17(tm790)* mutants have significantly impaired aversive associative learning and STAM performance compared to wild-type worms. Absolute values of learning and memory performance indices are reported. Mean ± SEM. n = 15 per genotype. ***p < 0.001, ****p < 0.0001; ns, not significant (p > 0.05). **(H)** Putative diagram of how INS-17 is upregulated in the nervous system to signal the valence of positive and aversive environmental conditions to inform behavioral outputs advantageous for survival.

### *ins-17* expression rapidly responds to nutrient deprivation

Next, we wanted to uncover the physiological conditions that regulate *ins-17* expression and assess their role in learning and memory. We mined publicly available gene expression datasets and discovered that *ins-17* transcription is likely regulated by metabolic state, as it is consistently upregulated in multiple starvation paradigms and genetic models of nutrient deprivation (S. Table 1) (49,51,59–62). Interestingly, our appetitive behavioral paradigm involves a 1-hour food deprivation (FD) step prior to the 1 CS-US (food-butanone) pairing, which has been reported to promote a more robust CS-US association compared to animals that do not undergo a FD period prior to memory training (31,63). We first examined if the FD step incorporated into our paradigm was sufficient to increase *ins-1*7. We performed qPCR analysis to measure *ins-17* levels in wild-type worms after 1 hour of FD and found that even this short time period was sufficient to observe a significant increase in *ins-17* expression (Fig. 5C, left). We also observed more robust upregulation of *ins-17* following 2 hours of FD (Fig. S5B), suggesting that this locus rapidly responds to changes in nutrient availability.

### INS-17 is required for properly sensing internal nutrient status to allow for processing of stimulus valence

Based on our gene expression results, we hypothesized that INS-17 could promote memory performance by functioning as a FD signal. We first assessed the role of insulin signaling in FD-associated improvements in associative behavior (Fig. 5C, right). We confirmed that FD could indeed improve memory ability by comparing the behavioral performance of wild-type animals that underwent FD prior to training to animals that were not food-deprived (non-FD) prior to training. Similar to previous reports (31), we found that wild-type FD animals have significantly better ITAM performance, which is measured one hour post-training, compared to non-FD animals (Fig. 5D, S5C-D). Interestingly, *daf-2* mutant phenotypes are strongly associated with phenotypes paralleling those of chronically starved worms (64–67), so we hypothesized that *daf-2* animals may not require the FD step prior to 1 CS-US pairing for robust ITAM behavior, as they are thought to have a perceived nutrient status of chronic starvation even when well-fed. We found that non-FD *daf-2* animals that underwent 1 CS-US pairing had ITAM behavior nearly identical to *daf-2* animals that were FD prior to training (Fig. 5E, S5E). Altogether, these results indicate that *daf-2* animals’ improved memory phenotype could be driven by their sensation of a chronically starved internal state.

We next tested if INS-17 could function as a FD signal. If this was the case, we hypothesized that increased levels of INS-17 could be sufficient to promote robust ITAM behavior in the absence of the FD step prior to 1 CS-US pairing. To test this, we examined if wild-type animals fed INS-17 peptide required FD for enhanced ITAM. We compared memory performance of SCR or INS-17 fed animals after training in the absence of FD, and found that, similar to *daf-2* mutants, INS-17 fed animals exhibited significantly more robust ITAM than SCR fed animals in the non-FD condition (Fig. 5F). Furthermore, FD had no detectable effect on ITAM performance in INS-17 fed animals, but significantly improved ITAM in SCR fed animals (Fig. S5F-I). These results suggest that INS-17 acts as a FD signal to promote a stronger association between a food and an odorant.

We next asked if INS-17 functions generally as a FD signal to promote associative behaviors using an aversive olfactory training paradigm, where an attractive odorant diacetyl is paired with the absence of food to form a negative association (68). If INS-17 functions as a signal of FD internal state, we hypothesized that *ins-17* mutants would have impaired aversive learning and memory compared to wild-type animals, because sensing starvation is required for a negative diacetyl association to be formed. Wild-type and *ins-17* loss-of-function mutants underwent well-established aversive memory assays using a diacetyl-starvation pairing (68). Strikingly, we found that *ins-17* is required for normal aversive associative behaviors, as *ins-17* animals had severe deficits in learning, STAM, and ITAM behavior compared to wild-type animals (Fig. 5G, S5J). The aversive and appetitive olfactory conditioning paradigms we tested here recruit distinct neuronal circuits (63,69–75). Therefore, our results suggest that *ins-17* may be broadly required in the nervous system to signal a FD state so that the valence of olfactory stimuli (positive when paired with food, negative when paired with starvation) can be processed appropriately to promote advantageous behaviors (Fig. 5H). This could explain how IIS acts in the nervous system to integrate metabolic status and regulate specific behavioral outputs, such as learning and memory.

## Discussion

Across species, insulin signaling is pleiotropic. In the worm, the DAF-2 receptor regulates a wide array of phenotypes, including those related to development, lifespan, metabolism, fecundity, cognitive behaviors, and motility, among others. How does a single receptor mediate such physiologically distinct phenotypes? Here we report that some of these pleiotropic actions result from specific ligand-receptor interactions. In this instance, we show that an olfactory memory phenotype is likely mediated by INS-17, an antagonistic ILP with no previously reported role in behavior (to our knowledge). DAF-2 receptor partial loss-of-function mutants have a well-established extended memory phenotype (31), so it is not surprising that a predicted antagonistic ILP regulated these behaviors. However, we were surprised that only a single strong antagonist appeared to be required in our paradigms. The importance of INS-17 to the extended memory phenotype is supported by our finding that overexpression through feeding is sufficient to recapitulate the memory phenotype of *daf-2* mutants. While we established that other antagonists are not required for this specific behavioral phenotype, many ILPs are diverse and act as agonists or antagonists in a context-dependent fashion, and most do not have a described role in associative behaviors. Future studies systemically characterizing these diverse ligands will be informative for defining DAF-2 receptor pleiotropy across tissues.

Interestingly, several transcriptomic datasets show that *ins-17* transcripts are increased roughly 2-fold in *daf-2* mutants and decreased in *daf-16* mutants compared to wild-type animals (Table 1) (34,49,51,59,60,62). This indicates that INS-17 is highly regulated by IIS signaling and is potentially acting both upstream and down-stream of DAF-2/IR. “ILP to ILP” signaling, where the action of one ILP affects the expression of another ILP, could be a component of *ins-17*’s role in integrating sensory cues and nutrient availability to elicit behavioral outputs. One example of “ILP to ILP” signaling in behavior is between INS-6 and INS-7 in aversive pathogen learning, where INS-6 released by the ASI neuron regulates *ins-7* transcription in the URX neurons to regulate this behavior (76). Further transcriptional analysis of *ins-17* mutants and INS-17 overexpression models could provide insight into which ILPs may be regulating or be regulated by INS-17.

We also report that INS-17’s function in adulthood is specific to behavior, as manipulating INS-17 does not affect lifespan (49). While *daf-2* animals’ extended lifespan is a canonical phenotype, there may exist alternate downstream IIS pathways that are dispensable for lifespan regulation. There are reports that the sole insulin receptor substrate homolog IST-1 may differentially activate distinct arms of the IIS signaling pathway, as it can function upstream of—or partially in parallel to—DAF-16 and is not required for normal growth and development (77–79). Thus, it is possible the 40 known ILPs may similarly engage the IIS pathway. In the present study, we investigated IST-1, which engages atypical IIS signaling to regulate aversive olfactory learning (79). Here, we report that *ist-1* mutants display positive olfactory memory deficits that were rescued by INS-17 peptide feeding; taken with our similar results in *daf-16* mutants, these findings suggest INS-17 may activate multiple arms of the IIS pathway to regulate adult cognitive behaviors. We find that *daf-16;ist-1* double mutants do not respond to INS-17 peptide supplementatio<u>n,</u> suggesting that, similar to other olfactory behavioral paradigms(79), INS-17 engages multiple pathways that can act partially in parallel to one another downstream of DAF-2 to regulate behavior. It will be intriguing in the future to examine if other DAF-2-mediated associative behaviors similarly engage both downstream pathways.

We also find that the genomic context in which an ILP is regulated provides specificity, as “swapping” a structurally similar insulin, *ins-37* (47), into *the ins-17* genetic locus is sufficient to functionally replace *ins-17* in behavior, though endogenous *ins-37* is not required for learning phenotypes in our paradigm. This result supports that tissue-specific gene regulation may inform DAF-2/IR pleiotropy, warranting future studies exploring tissue-specific requirements for DAF-2/IR function.

Furthermore, we demonstrate that *ins-17* is required for both positive and aversive associative behaviors. Both associative behavioral paradigms include a relatively short, 1-hour FD step prior to training to prime animals to form a stronger association between food and an odorant, thus promoting a more robust behavioral response after training. This FD step occurs within the timeframe before animals display significant changes in sensory integration behavior due to a “lack of food sensation”, which begins after 3 hours of food-deprivation (80); however, food is cleared from the alimentary system within 2 minutes of ingestion (81,82), suggesting the need for a FD signal to inform robust behavioral responses after only 1 hour of FD. Our gene expression data, similar to previous studies (51,59,60), shows that the *ins-17* gene is activated in this timeframe, suggesting that could it be involved in responding to these brief FD signals. This notion is supported by our findings that feeding INS-17 to animals that do not undergo FD results in normal learning and memory, bypassing the need for FD to promote memory. Together, these results suggest that INS-17 could be released in response to acute and prolonged FD to inform sensory integration behavior. Interestingly, the only phenotype previously associated with INS-17 is dauer formation to protect progeny from unfavorable conditions (49), namely starvation (83). We speculate that INS-17 may respond to FD to coordinate phenotypic responses in different life stages that are most advantageous; for example, while *ins-17* regulates dauer formation, which is a response to lack of food availability and other stresses during development, it may also regulate the ability to form associations in the context of food availability in adulthood, which is beneficial for egg-laying and progeny survival (64,84). In the future, it will be interesting to further examine how both the *ins-17* locus and INS-17 secretion are regulated in the context of changing nutrient availability.

Since *ins-17* is required for both positive and aversive associative behaviors, our results support that INS-17 may function as a broad signal for nutrient deprivation to the nervous system. *ins-17* is widely expressed across the nervous system (58,61), but is most highly expressed in the AIZ interneuron, which mediates a variety of behavioral outputs in response to multiple sensory neurons and circuits (85–87). We observed that associative behavior impairments resulting from loss of ins-17 do not stem from baseline sensory deficits, which strongly indicates that rather sensory information may not be integrated correctly or that sensory neurons cannot alter their responses to an environmental stimulus in the context of altered metabolic status. We have established that INS-17 may be essential for interpreting internal state and proper valence processing. In mammals, neuropeptides also are important for signaling valence in positive and aversive associative paradigms (88,89), and this is likely a theme of peptides modulating circuits more broadly. Future studies examining how INS-17 regulates sensory circuit activity in various contexts will help further define *daf-2* pleiotropic mechanisms.

Our results bring an attractive model for the expansion of the ILP family in *C. elegans*. ILPs are encoded across the genome and function under the control of a variety of regulatory elements so the animal can dynamically respond to a combination of nutrients and environmental stimuli via DAF-2/IR to facilitate the most advantageous phenotype. Having a diverse array of peptides that can carry out distinct outputs is beneficial, especially if there is only one receptor to mediate such outputs. In the case of INS-17, being able to properly make associations in the face of changing environments can promote survival, such as remembering availability (or lack) of a food source associated with specific odors. Together, our findings have identified INS-17 as a physiological signal that is required for positive and aversive associative behaviors and may integrate environmental cues and nutrient status to inform valence processing and regulate behavioral outcomes. This research underscores the importance of future work investigating the rich and diverse network of DAF-2/IR signaling.

## Methods

### General *C. elegans* maintenance

Worms were maintained at 20 °C on plates containing either 1) Standard nematode growth medium (NGM): 3 g/L NaCl, 17 g/L agar, 2.5 g/L peptone, 25 mL/L KPO4 (pH 6.0), 1 mL/L MgSO4, 1 mL/L CaCl2, and 1 mL/L cholesterol (5 mg/mL in ethanol) (90); or 2) High growth medium (HGM): 3 g/L NaCl, 30 g/L agar, 20 g/L peptone, 25 mL/L KPO4 (pH 6.0), 1 mL/L MgSO4, 1 mL/L CaCl2, and 4 mL/L cholesterol (5 mg/mL in ethanol) (50). Plates were seeded with an *E. coli* OP50 lawn for ad libitum feeding (90).

Animals were developmentally synchronized using an alkaline-bleach solution (1.5 mL 5 N NaOH, 3.0 mL 5% sodium hypochlorite, 5.5 mL water) to collect eggs from gravid adults; eggs were subsequently washed three times with M9 buffer solution (6 g/L Na2HPO4, 3 g/L KH2PO4, 5 g/L NaCl and 1 mL/L 1M MgSO4 in ultrapure water) before plating (90).

### Strains

Wild-type: N2 Bristol

Mutants: *ins-17(tm790), ins-21(tm5180)*, *ins-37(tm6892)*, and *ins-39(tm6520)* were obtained from the National BioResource Project, Japan.

RB2489 (*ins-15(ok3444)*), RB2594 (*ins-22(ok3616)*), CB1370 (*daf-2(e1370)*), CF1038 (*daf-16(mu86)*), MAH677 (*sid-1(qt9) V; sqIs71[rgef-2p::GFP + rgef-1p::sid-1]*) were obtained from the Caenorhabditis Genetics Center.

CX17790 (*ist-1(ok2706)* was described previously(79) and generously gifted to us by Dr. Cornelia Bargmann.

Strains made in collaboration with InVivo Biosystems include *ins-17* CRISPR KO animals with *ins-37* replacement COP2848 (*ins-17(knu1330[ins-37]) III*) and *ins-17* CRISPR KO animals COP2835 (*ins-17(knu1320) III*).

The following strains were generated by crosses: RNA34 (*daf-2(e1370)*; *ins-17(tm790))* and RNA59 (*daf-16(mu86); ist-1(ok2706)*).

### Peptide plasmid generation and transformation

The amino acid coding sequence for each peptide was identified using www.wormbase.org (57). These sequences were codon-optimized for *C. elegans* and cloned in pET-21a(+) cloning vector via XhoI/XhoI strategy by GenScript. The scramble control sequence and plasmid were generated using the same cloning strategy by GenScript. Plasmids arrived lyophilized and were re-suspended in nuclease-free H_2_O to a concentration of 100ng/μl and stored at −20 °C. Each expression clone was transformed into competent iOP50 cells using methods previously described(91). Briefly, iOP50 stocks were streaked out onto LB plates containing 50 μg/mL tetracycline and grown at 37 °C overnight. Then, single colonies of iOP50 were isolated and used to inoculate 2.0 mL LB containing 50 μg/ml tetracycline and placed in a 37 °C shaker overnight. 0.5 mL of each culture was used to inoculate 3.0 mL of LB without antibiotics and grown for 2 hours in a 37 °C shaker. Each culture was then aliquoted in 0.75 mL increments into four 1.5 mL tubes and cooled for 10 minutes on ice. Next, the cells were harvested by centrifugation at 6,000 rpm for 5 minutes at 4 °C. The resulting supernatant was discarded, and cells were placed back on ice and re-suspended in 1.0 mL 100 mM ice-cold CaCl_2_ solution. Cells were kept on ice for 20 minutes and then harvested by centrifugation as in the previous step. After discarding the supernatant, cells were re-suspended in 150μl of ice-cold CaCl_2_ solution and placed on ice to be immediately used for transformation.

Competent cells were transformed using the New England Biolabs protocol (C2987H/C2987I). Single colonies of transformed cells were isolated and cultured in 3.0 mL of LB containing 50 μg/mL tetracycline and 50 μg/mL carbenicillin overnight in a 37 °C shaker. Cultures were frozen down as glycerol stocks and stored at −80 °C. Successful transformation was verified by isolating plasmids from cultures using a Zyppy Plasmid Miniprep Kit and sending samples for whole-plasmid sequencing (Plasmidsaurus).

### Peptide plate preparation

LB containing 50 μg/mL tetracycline and 50 μg/mL carbenicillin was inoculated with iOP50 encoding either the peptide sequence of interest or scramble peptide sequence and cultured for 18-20 hours in a 37 °C shaker. 100 mm NGM plates containing 50 μg/mL tetracycline, 50 μg/mL carbenicillin, and 1.0 mL/L isopropyl-β-D-thiogalactoside (IPTG) were plated with 1.0 mL of corresponding culture. Seeded plates were left to dry at least 24 hours at room temperature and never kept longer than 72 hours before use. Approximately one hour before transferring worms to peptide or scramble control plates, 200 μL of 0.1 M IPTG was added to each plate and left to dry.

### Peptide feeding

We adapted a previously published peptide feeding-based approach (53). Worms were maintained on NGM agar plates seeded with OP50 *E. coli* and maintained at 20 °C until the start of experiments. All animals were transferred to a conical tube and washed three times with M9 buffer solution before being transferred to NGM plates containing 50 μg/mL Tetracycline, 50 μg/mL Carbenicillin, and 1.0 mM IPTG that were seeded with 1.0 mL of iOP50 expressing either the scramble peptide or the peptide of interest. Animals were kept on the scramble or peptide-expressing lawns for at least 48 hours of exposure time at 20 °C before performing assays. Exact peptide feeding timeline parameters are expounded in the results section.

### RNAi treatment

Worms were maintained on NGM agar plates seeded with OP50 *E. coli* at 20 °C until the start of experiments. At the L4 stage, animals were transferred to a conical tube and washed three times with M9 buffer solution before being transferred to NGM plates containing 50 μg/mL Carbenicillin and 1.0 mM IPTG that were seeded with 1.0 mL of HT115 bacteria expressing either RNAi against the gene of interest or the vector control RNAi. Animals were kept on the RNAi lawns for 48 hours of exposure time at 20 °C before performing assays at Day 2 of adulthood.

### Naïve chemotaxis assay

Chemotaxis assays were performed using a protocol based on previously published assays(92). Briefly, assays were performed on unseeded 100mm NGM plates. On the back of each plate, two marks were made on opposite sides of the plate, approximately 5mm from the edge. 1μL of sodium azide (Thermo Fisher) was placed on both spots and allowed to dry before adding 1μL of odorant (Sigma Aldrich) diluted in ethanol on one mark and 1μL ethanol on the other. Using M9 buffer, worms were washed off their plates and subsequently washed three times to eliminate any leftover bacteria that could impact baseline behavior. Then, worms were placed near the bottom center of the plate, equidistant between the two marks, and allowed to chemotax for an hour. Chemotaxis indices were calculated as follows:

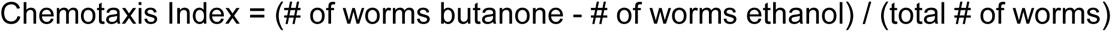

For naïve chemotaxis assays performed to assess baseline chemosensory function, we tested the naïve preference to a highly appetitive concentration of butanone (0.1%) as previously reported (92).

### Positive associative butanone learning and memory assay

Worms were trained and tested for learning, short- and intermediate-term memory changes as previously described (50). Briefly, synchronized adult worms were washed off plates with M9 buffer and allowed to settle by gravity, followed by two subsequent washes with M9 buffer to remove leftover bacteria that could alter baseline behavior. Next, the worms were food-deprived for one hour in M9 buffer before exposure to 1 CS-US pairing, where food-deprived worms were transferred to 100mm NGM conditioning plates seeded with OP50 *E*. *coli* bacteria (or iOP50 bacteria containing a peptide or scramble control plasmid for peptide feeding experiments) and had 16μL of 10% butanone (Sigma Aldrich) diluted in ethanol streaked on the lid in a ‘#’ shape for one hour. After conditioning, the trained population of worms were tested for chemotaxis to 10% butanone and to an ethanol control using standard, previously described chemotaxis conditions(92). Different stages of memory were tested by measuring chemotaxis indices of separate populations of worms at timepoints including 0 min. (learning), 30 min. (STAM), 60 min. (ITAM) and 120 min. (forgetting) after training. For timepoints after learning behavior, worms were transferred to hold plates seeded with OP50 *E. coli* or iOP50 bacteria for peptide feeding experiments as stated above.

Chemotaxis indices for each timepoint were calculated as detailed in “Naïve chemotaxis assays” section.

Performance index is the change in the chemotaxis index after training relative to the naïve (untrained) chemotaxis index:

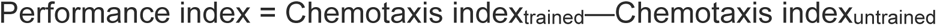

% Maximum Performance Indices are performance indices represented as a percentage of the maximum value that can be performed relative to the naïve chemotaxis index(31,36). This % can be more informative for visualizing performance indices when naïve chemotaxis behavior is high, as in *daf-2(e1370)* animals. This is calculated for each timepoint by dividing the performance index of a timepoint by the average of the untrained chemotaxis indices of the same behavioral replicate subtracted from 1:

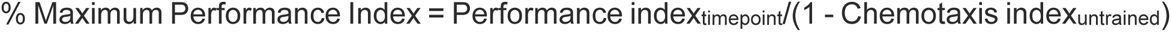

### Butanone enhancement assay

This assay was adapted from previously described methods (63). Unlike the positive butanone associative memory assays, butanone enhancement assays skip the 1-hour food-deprivation step prior to the conditioning step. All other methods are identical to those listed in the positive butanone associative memory assay section.

### Negative associative learning and memory assay

Worms were tested for learning and memory changes in response to aversive diacetyl training, adapted from previously published methods (68). Similar to positive associative assays, synchronized Day 1 adult worms were washed off plates with M9 buffer and then allowed to settle by gravity and washed twice more with M9 buffer. After washing, the worms were conditioned for one hour by food-deprivation on 100mm NGM plates with no bacteria and 16uL of 100% diacetyl (Sigma Aldrich) streaked on the lid in a ‘#’ shape. The trained population of worms were then tested for chemotaxis to 1% diacetyl and to an ethanol control using standard, previously described chemotaxis conditions (92). Learning and memory performance indices were calculated for each timepoint as previously stated in the positive butanone associative assay section.

### Lifespan assay

Lifespan assays were performed on *ins-17(tm790)* animals and wild-type animals that were bleached onto NGM plates seeded with OP50 *E. coli* and maintained at 20 °C; at the L4 larval stage, worms were picked for survival analysis and transferred every other day to freshly seeded plates. Similarly, lifespan assays with peptide feeding administration were performed on wild-type animals that were bleached onto NGM plates seeded with OP50 *E. coli* and maintained at 20 °C. Then, at the L4 larval stage, worms were picked for survival analysis and transferred every other day to NGM plates containing 50 μg/mL tetracycline, 50 μg/mL carbenicillin, and 1.0 mL/L IPTG that were seeded with either INS-17 iOP50 or scramble iOP50 *E. coli*. Lifespans began (t = 0) at Day 1 of adulthood and had an n = ∼120 worms per strain/trial for a total of three biological replicates per assay.

### Food-deprivation behavioral experiments

For food-deprivation experiments, well-fed, synchronized adult worms were washed off plates using M9 buffer into a 15mL conical and washed two more times to remove bacteria that could alter baseline behavior. Next, worms were either 1) food-deprived for one hour on a 100mm NGM plate without food or 2) non-food deprived for one hour by being transferred to a 100mm plate seeded with a bacterial lawn (iOP50 bacteria for peptide feeding experiments). Worms were then immediately exposed to the conditioning step (1 CS-US pairing) and stages of learning and memory were tested as stated in the positive associative butanone learning and memory assays section.

### RNA isolation, cDNA synthesis and qRT-PCR

For qRT-PCR food-deprivation experiments, well-fed, synchronized adult worms were washed off plates using M9 buffer into a 15mL conical and washed two more times to remove bacteria that could alter baseline behavior. Next, worms were either 1) food-deprived for one hour or two hours after being transferred to a 100mm plate with no bacterial lawn or 2) non-food deprived for one hour or two hours after being transferred to a 100mm plate seeded with a bacterial lawn.

After feeding or food deprivation, worms were crushed in liquid nitrogen using a mortar and pestle and added to Trizol (Thermo Fisher Scientific). RNA was isolated per manufacturer’s instructions, followed by DNase treatment (Qiagen). cDNA was synthesized with an oligo dT primer and Superscript III reverse transcriptase enzyme (Thermo Fisher Scientific). cDNA was mixed with buffers, primers, SYBR green, and hot start Taq polymerase in a master mix prepared by a manufacturer (Thermo Fisher Scientific). PCR reactions were run using a Quant Studio 7 Pro Dx Real-Time PCR System (Thermo Fisher Scientific), followed by a dissociation reaction to determine specificity of the amplified product. Gene expression was quantified using the ΔΔCt method using *pmp-3* as a reference gene. Primer sets were as follows:

*ins-17* For: 5’-GGACACTATCCGACCACCAC -3’:
*ins-17* Rev: 5’-TTCACATCCCGTCTCACAGC -3’:
*pmp-3* For: 5’-AGTTCCGGTTGGATTGGTCC -3’
*pmp-3* Rev: 5’-CCAGCACGATAGAAGGCGAT-3’

### Statistical analysis and software

All statistics (excluding survival analysis) were calculated with GraphPad Prism Software. All statistical data are reported in the main text, figures, and tables. Significance threshold of p < 0.05 was used. The symbols *, **, ***, and **** refer to p < 0.05, 0.01, 0.001, and 0.0001. The symbols #, ##, ###, and #### refer to p < 0.05, 0.01, 0.001, and 0.0001. For comparing performance indices between two behavior conditions (e.g. *vector control* vs *ins-17 RNAi*), we used a Mann-Whitney test comparing ranks, as it does not assume normality. For comparing performance indices between three or more groups (e.g. wild-type vs multiple insulin receptor antagonist mutants), we performed one-way analysis of variances followed by Bonferroni post hoc tests for multiple comparisons. Two-way ANOVAs were used for evaluating effects between genotype and timepoint (T0, T30, T60, T120) on performance indices with a significant interaction between factors (p<0.0001) that prompted Bonferroni post-hoc analyses to determine differences between individual groups. Sample size n represents the number of chemotaxis assays performed for behavioral experiments, with each assay containing approximately 50–150 worms. All experiments were performed on different days with distinct populations to confirm reproducibility. For qPCR analysis, unpaired t-tests with Welch’s corrections were performed to determine significance between conditions. Survival analysis statistics were generated using the log-rank (Mantel-Cox) method to test the null hypothesis in Kaplan-Meier survival analysis and evaluated using OASIS survival analysis software (93).

## Acknowledgements

We thank the Bargmann lab for generously sharing the *ist-1(ok2706)* strain, and the Caenorhabditis Genetics Center for most other strains used in this publication. We also thank Ben Jussila and In Vivo Biosystems for strains and helpful discussion. Additionally, we especially thank Drs. Buck Samuel and Shinya Yamamoto at Baylor College of Medicine for helpful discussion and advice at various stages of the project. Finally, we thank Arey lab members Katie L. Brandel-Ankrapp and Ty M. Gadberry for feedback on the manuscript.

