## Supplemental Figures, Legends, and Table for "INS-17, an insulin-receptor ligand, confers specificity in adult IIS-regulated phenotypes in *C. elegans*" (from another version of this preprint: version 1, 2025-11-21)

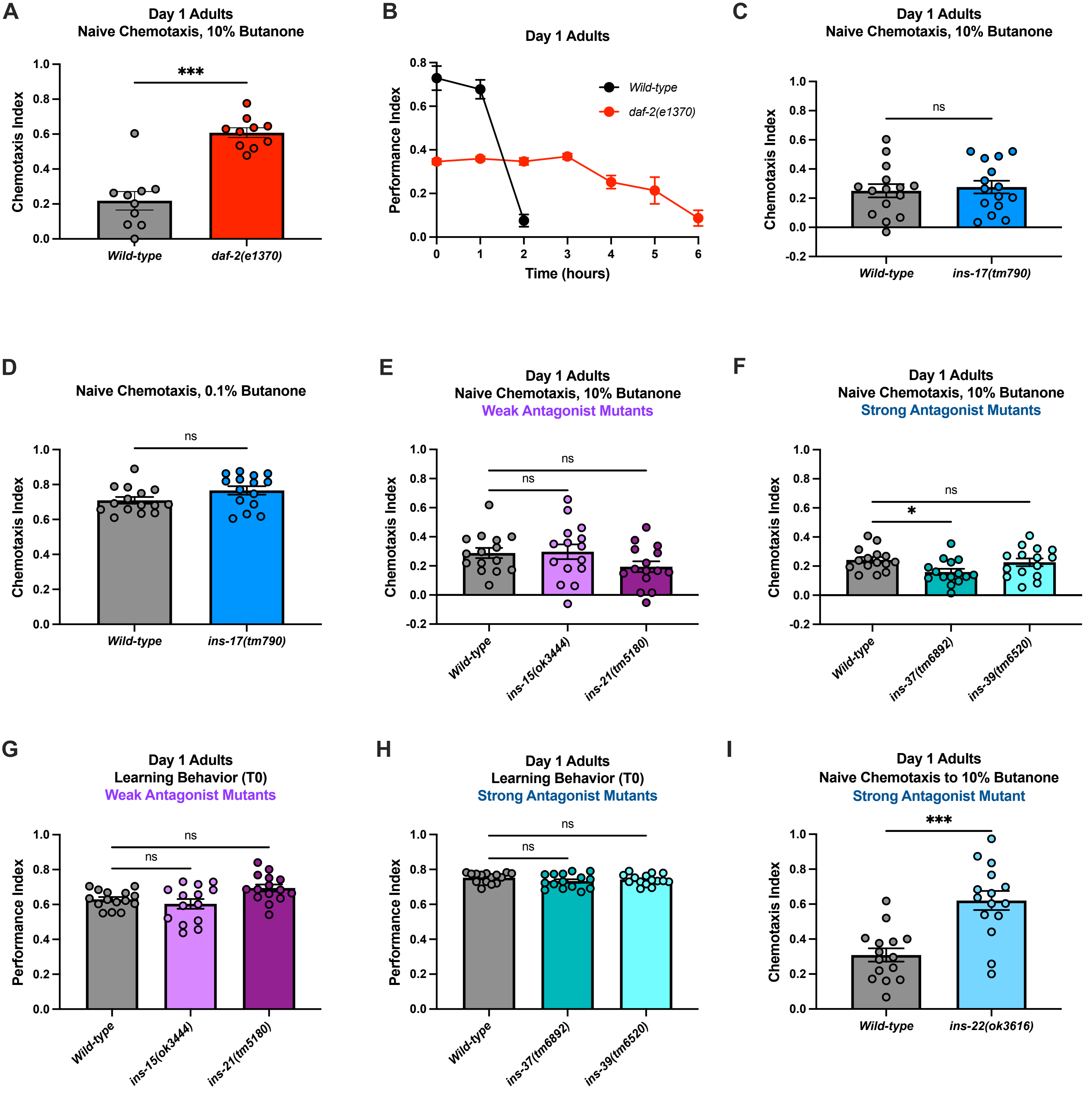
A
Day 1 Adults
Naive Chemotaxis, 10% Butanone
Chemotaxis Index
***
Wild-type
daf-2(e1370)
B
Day 1 Adults
Performance Index
Wild-type
daf-2(e1370)
Time (hours)
C
Day 1 Adults
Naive Chemotaxis, 10% Butanone
Chemotaxis Index
ns
Wild-type
ins-17(tm790)
D
Naive Chemotaxis, 0.1% Butanone
Chemotaxis Index
ns
Wild-type
ins-17(tm790)
E
Day 1 Adults
Naive Chemotaxis, 10% Butanone
Weak Antagonist Mutants
Chemotaxis Index
ns
ns
Wild-type
ins-15(ok3444)
ins-21(tm5180)
F
Day 1 Adults
Naive Chemotaxis, 10% Butanone
Strong Antagonist Mutants
Chemotaxis Index
ns
*
Wild-type
ins-37(tm6892)
ins-39(tm6520)
G
Day 1 Adults
Learning Behavior (T0)
Weak Antagonist Mutants
Performance Index
ns
ns
Wild-type
ins-15(ok3444)
ins-21(tm5180)
H
Day 1 Adults
Learning Behavior (T0)
Strong Antagonist Mutants
Performance Index
ns
ns
Wild-type
ins-37(tm6892)
ins-39(tm6520)
I
Day 1 Adults
Naive Chemotaxis to 10% Butanone
Strong Antagonist Mutant
Chemotaxis Index
***
Wild-type
ins-22(ok3616)

**Figure S1**

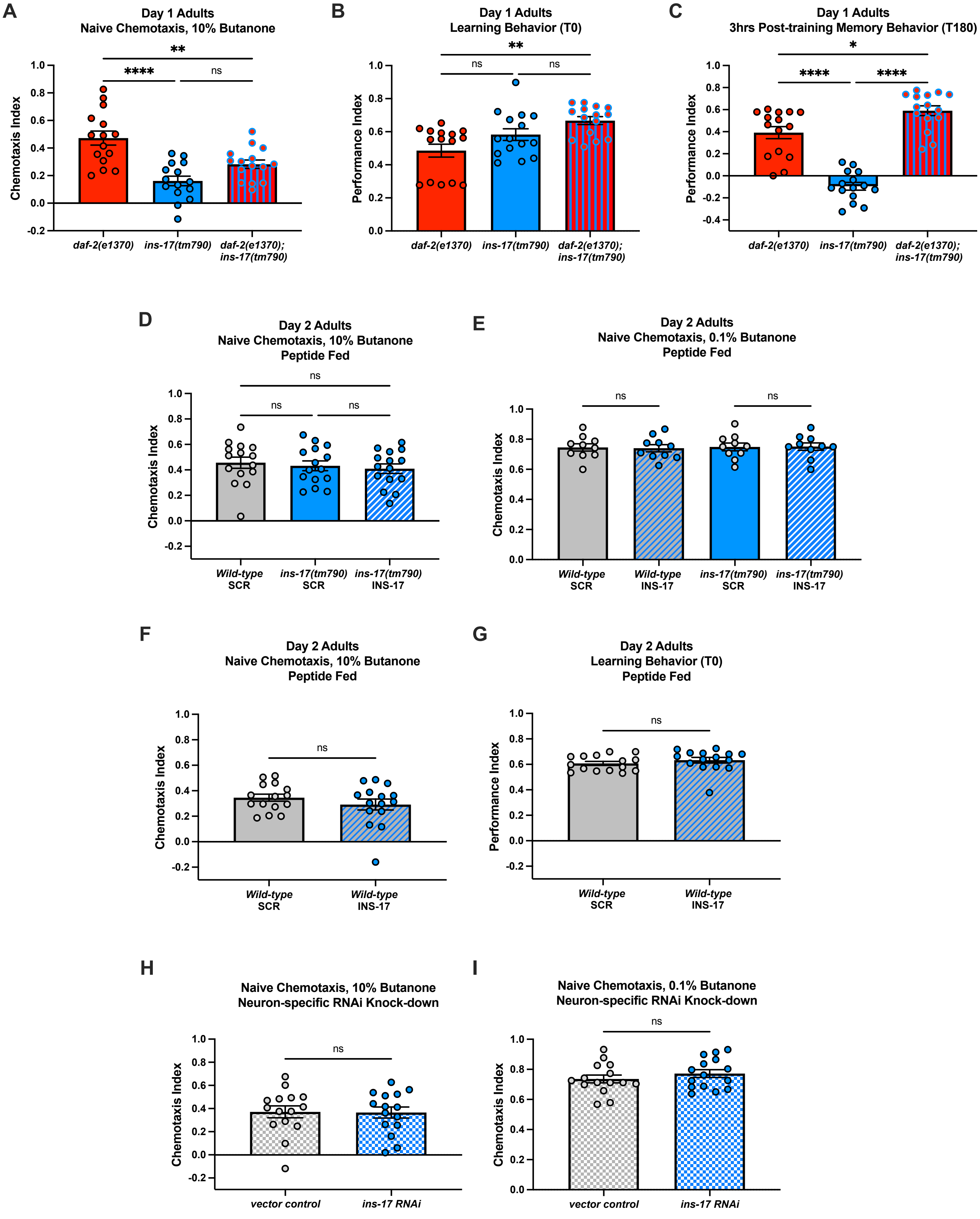
A
Day 1 Adults
Naive Chemotaxis, 10% Butanone
Chemotaxis Index
daf-2(e1370)
ins-17(tm790)
daf-2(e1370); ins-17(tm790)
B
Day 1 Adults
Learning Behavior (T0)
Performance Index
C
Day 1 Adults
3hrs Post-training Memory Behavior (T180)
Performance Index
D
Day 2 Adults
Naive Chemotaxis, 10% Butanone
Peptide Fed
Wild-type SCR
ins-17(tm790) SCR
ins-17(tm790) INS-17
E
Day 2 Adults
Naive Chemotaxis, 0.1% Butanone
Peptide Fed
Wild-type SCR
Wild-type INS-17
ins-17(tm790) SCR
ins-17(tm790) INS-17
F
Day 2 Adults
Naive Chemotaxis, 10% Butanone
Peptide Fed
G
Day 2 Adults
Learning Behavior (T0)
Peptide Fed
H
Naive Chemotaxis, 10% Butanone
Neuron-specific RNAi Knock-down
vector control
ins-17 RNAi
I
Naive Chemotaxis, 0.1% Butanone
Neuron-specific RNAi Knock-down

Figure S2

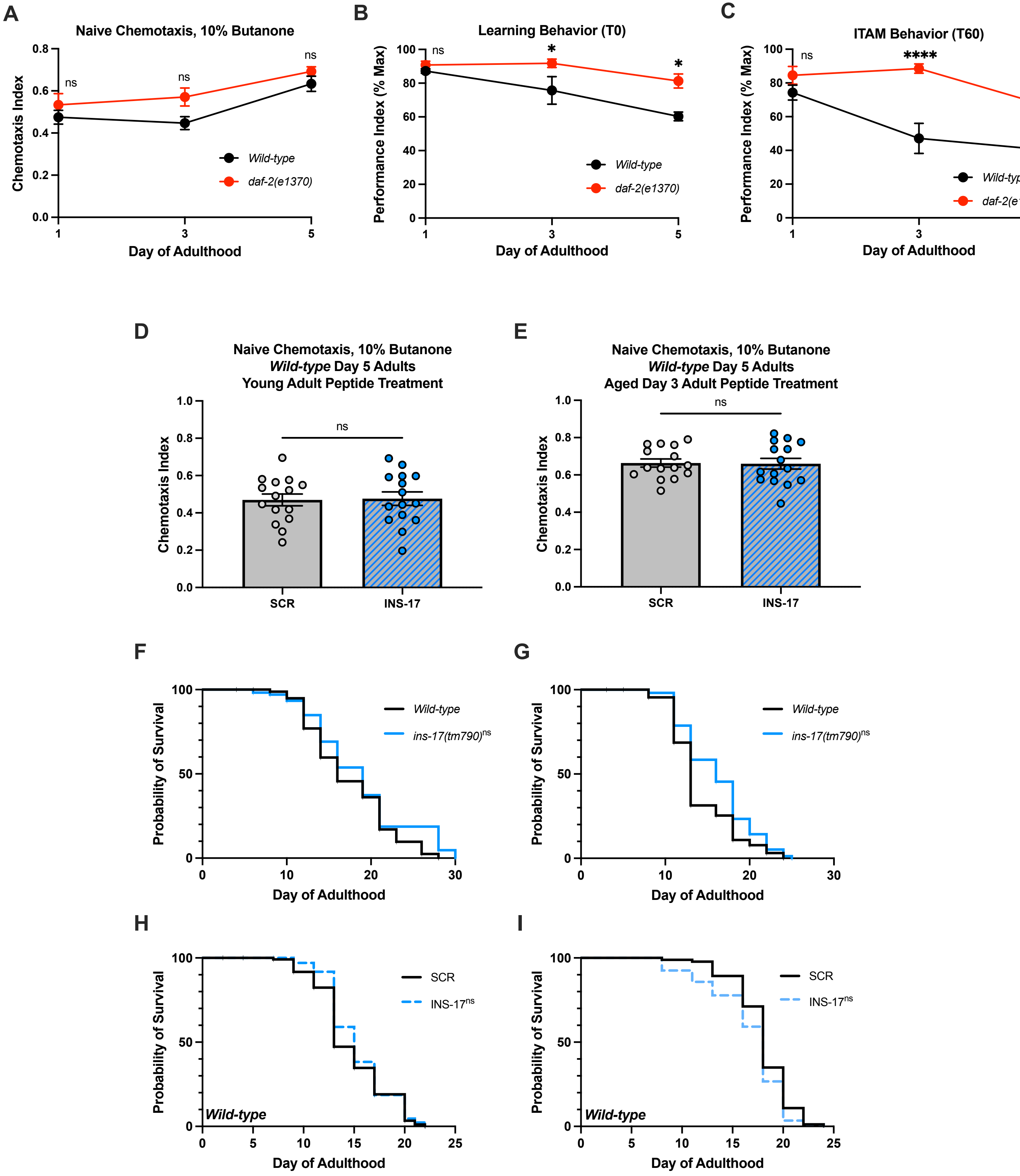

A
Naive Chemotaxis, 10% Butanone
Chemotaxis Index
Day of Adulthood
Wild-type
daf-2(e1370)
ns
B
Learning Behavior (T0)
Performance Index (% Max)
Day of Adulthood
Wild-type
daf-2(e1370)
C
ITAM Behavior (T60)
Performance Index (% Max)
Day of Adulthood
Wild-type
daf-2(e1370)
D
Naive Chemotaxis, 10% Butanone
Wild-type Day 5 Adults
Young Adult Peptide Treatment
Chemotaxis Index
SCR
INS-17
E
Naive Chemotaxis, 10% Butanone
Wild-type Day 5 Adults
Aged Day 3 Adult Peptide Treatment
Chemotaxis Index
SCR
INS-17
F
Probability of Survival
Day of Adulthood
Wild-type
ins-17(tm790)ns
G
Probability of Survival
Day of Adulthood
Wild-type
ins-17(tm790)ns
H
Probability of Survival
Day of Adulthood
SCR
INS-17ns
Wild-type
I
Probability of Survival
Day of Adulthood
SCR
INS-17ns
Wild-type


**Figure S3**

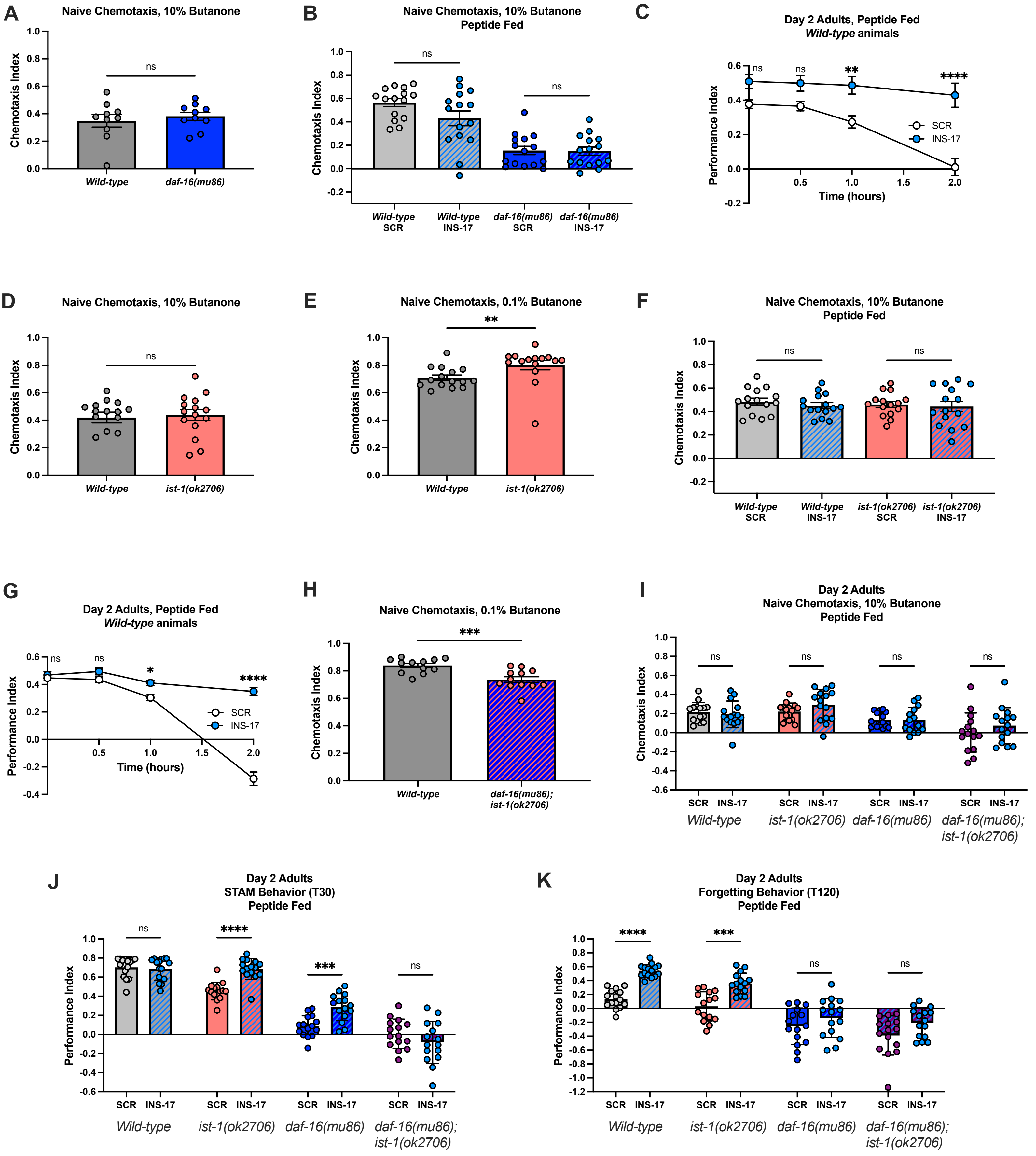
A
Naive Chemotaxis, 10% Butanone
Chemotaxis Index
ns
Wild-type
daf-16(mu86)
B
Naive Chemotaxis, 10% Butanone
Peptide Fed
Chemotaxis Index
ns
ns
Wild-type SCR
Wild-type INS-17
daf-16(mu86) SCR
daf-16(mu86) INS-17
C
Day 2 Adults, Peptide Fed
Wild-type animals
Performance Index
ns
ns
**
****
SCR
INS-17
Time (hours)
D
Naive Chemotaxis, 10% Butanone
Chemotaxis Index
ns
Wild-type
ist-1(ok2706)
E
Naive Chemotaxis, 0.1% Butanone
Chemotaxis Index
**
Wild-type
ist-1(ok2706)
F
Naive Chemotaxis, 10% Butanone
Peptide Fed
Chemotaxis Index
ns
ns
Wild-type SCR
Wild-type INS-17
ist-1(ok2706) SCR
ist-1(ok2706) INS-17
G
Day 2 Adults, Peptide Fed
Wild-type animals
Performance Index
ns
ns
*
****
SCR
INS-17
Time (hours)
H
Naive Chemotaxis, 0.1% Butanone
Chemotaxis Index
***
Wild-type
daf-16(mu86); ist-1(ok2706)
I
Day 2 Adults
Naive Chemotaxis, 10% Butanone
Peptide Fed
Chemotaxis Index
ns
ns
ns
ns
SCR
INS-17
Wild-type
ist-1(ok2706)
daf-16(mu86)
daf-16(mu86); ist-1(ok2706)
J
Day 2 Adults
STAM Behavior (T30)
Peptide Fed
Performance Index
ns
****
***
ns
SCR
INS-17
Wild-type
ist-1(ok2706)
daf-16(mu86)
daf-16(mu86); ist-1(ok2706)
K
Day 2 Adults
Forgetting Behavior (T120)
Peptide Fed
Performance Index
****
***
ns
ns
SCR
INS-17
Wild-type
ist-1(ok2706)
daf-16(mu86)
daf-16(mu86); ist-1(ok2706)

**Figure S4**

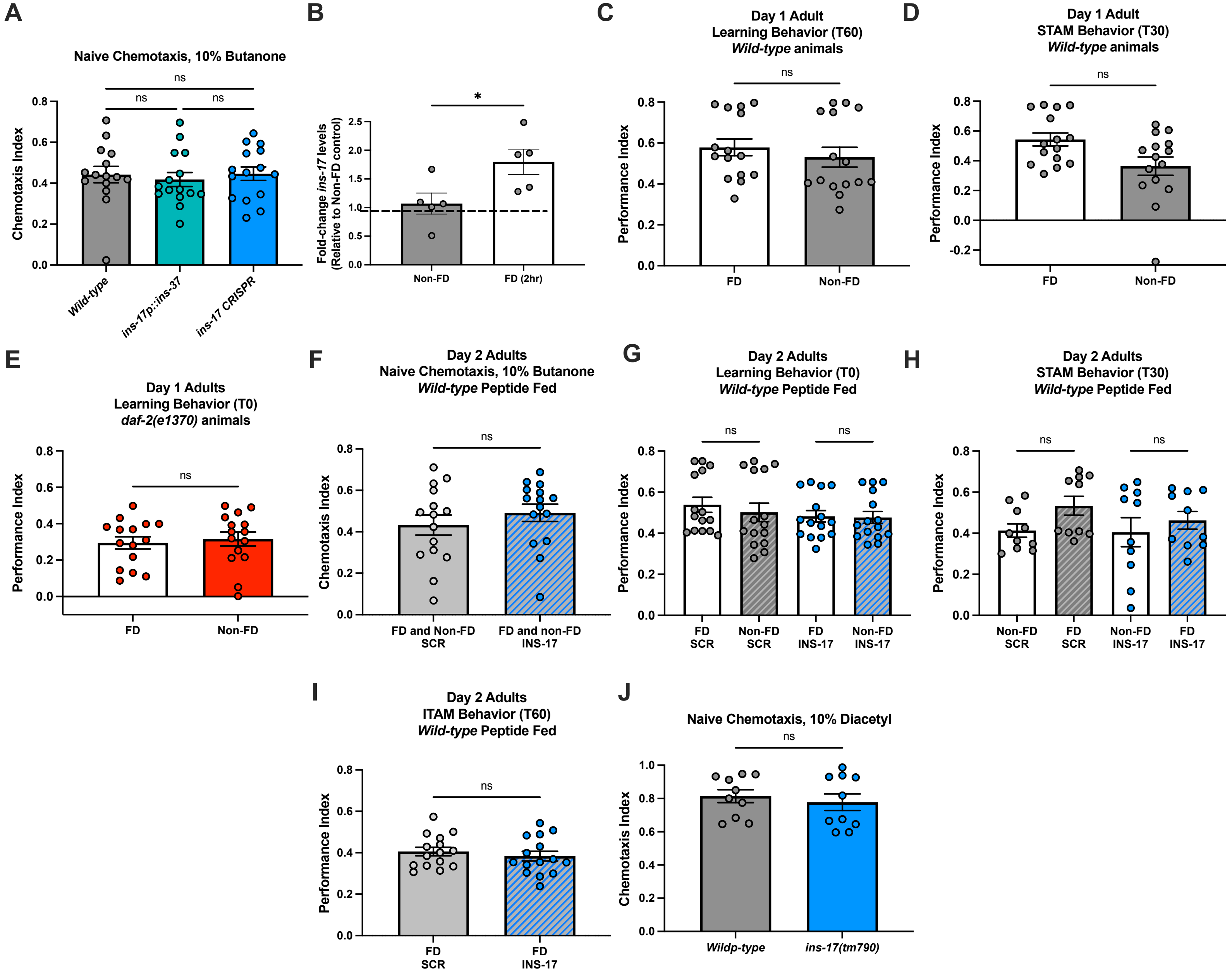
A
Naive Chemotaxis, 10% Butanone
ns
ns
ns
Chemotaxis Index
0.8
0.6
0.4
0.2
0.0
Wild-type
ins-17p::ins-37
ins-17 CRISPR
B
*
Fold-change ins-17 levels (Relative to Non-FD control)
2.5
2.0
1.5
1.0
0.5
0.0
Non-FD
FD (2hr)
C
Day 1 Adult
Learning Behavior (T60)
Wild-type animals
ns
Performance Index
0.8
0.6
0.4
0.2
0.0
FD
Non-FD
D
Day 1 Adult
STAM Behavior (T30)
Wild-type animals
ns
Performance Index
0.8
0.6
0.4
0.2
0.0
-0.2
FD
Non-FD
E
Day 1 Adults
Learning Behavior (T0)
daf-2(e1370) animals
ns
Performance Index
0.8
0.6
0.4
0.2
0.0
FD
Non-FD
F
Day 2 Adults
Naive Chemotaxis, 10% Butanone
Wild-type Peptide Fed
ns
Chemotaxis Index
0.8
0.6
0.4
0.2
0.0
FD and Non-FD SCR
FD and non-FD INS-17
G
Day 2 Adults
Learning Behavior (T0)
Wild-type Peptide Fed
ns
ns
Performance Index
0.8
0.6
0.4
0.2
0.0
FD SCR
Non-FD SCR
FD INS-17
Non-FD INS-17
H
Day 2 Adults
STAM Behavior (T30)
Wild-type Peptide Fed
ns
ns
Performance Index
0.8
0.6
0.4
0.2
0.0
Non-FD SCR
FD SCR
Non-FD INS-17
FD INS-17
I
Day 2 Adults
ITAM Behavior (T60)
Wild-type Peptide Fed
ns
Performance Index
0.8
0.6
0.4
0.2
0.0
FD SCR
FD INS-17
J
Naive Chemotaxis, 10% Diacetyl
ns
Chemotaxis Index
1.0
0.8
0.6
0.4
0.2
0.0
Wildp-type
ins-17(tm790)

**Figure S5**

**S1: *daf-2(e1370)* animals and *ins-22(ok3616)* animals have higher baseline naïve chemotactic preference for 10% butanone.**

**(A)** *daf-2(e1370)* animals have a higher baseline naïve chemotaxis to 10% butanone compared to wild-type animals. **(B)** Raw performance indices for wild-type animals decline at two hours compared to *daf-2(e1370)* mutants at six hours after 1 CS-US pairing. Mean ± SEM. n = 10 per genotype. ***p < 0.001. **(C)** *ins-17(tm790)* animals have no detectable impairments in baseline naïve chemotaxis to the neutral concentration of butanone (10%) **(D)** nor the appetitive concentration of butanone (0.1%) compared to wild-type. Mean ± SEM. n = 15 per genotype. ns, not significant (p > 0.05). **(E)** Weak (*ins-15, ins-21*) and **(F)** strong (*ins-39*) insulin receptor antagonist mutants have no detectable deficits in baseline naïve chemotaxis to 10% butanone. Strong antagonist *ins-37* has a slight impairment compared to wild-type, but this deficit does not appear to affect learning and memory behaviors. Mean ± SEM. n = 15 per genotype. *p < 0.05; ns, not significant (p > 0.05). **(G)** Weak and **(H)** strong antagonist mutants have normal learning behavior compared to wild-type after 1 CS-US pairing. Mean ± SEM. n = 15 per genotype. ns, not significant (p > 0.05). **(I)** *ins-22(ok3616)* animals have higher naïve attraction to 10% butanone that impair the ability to measure meaningful deficits in learning and memory behaviors. Mean ± SEM. n = 15 per genotype. ***p < 0.001.

**S2: Peptide administration (SCR and INS-17) and RNAi feeding (vector and *ins-17* RNAi) do not affect baseline naïve attraction to 10% butanone.**

**(A)** *daf-2(e1370)* animals have a higher baseline naïve chemotaxis to 10% butanone compared to *ins-17(tm790)* and *daf-2(e1370)*; *ins-17(tm790)* animals. This higher naïve preference appears to contribute to **(B)** *daf-2(e1370)* learning and **(C)** 3hr-memory deficits compared to *daf-2(e1370)*; *ins-17(tm790)* double mutants, but we find this significance is lost when comparing % Max Performance Indices as in **Fig. 2A**. Mean ± SEM. n = 15 per genotype. *p < 0.05, **p < 0.01, ****p < 0.0001; ns, not significant (p > 0.05). **(D)** SCR and INS-17 peptide treatments have no detectable effects on naïve chemotaxis behavior of wild-type worms nor *ins-17* mutants to 10% butanone as well as **(E)** 0.1% butanone. Mean ± SEM. n = 10 - 15 per genotype. ns, not significant (p > 0.05). **(F)** Naïve chemotaxis and **(G)** learning performance indices corresponding to **Fig. 2D.** Mean ± SEM. n = 15 per genotype. ns, not significant (p > 0.05). **(H)** Neuron-specific RNAi treatment has no detectable effect on naïve chemotaxis to 10% butanone nor **(I)** 0.1% butanone. Mean ± SEM. n = 15 per genotype. ns, not significant (p > 0.05).

**S3: *ins-17(tm790)* animals and manipulation of INS-17 levels via peptide administration have no detectable changes in lifespan compared to wild-type animals and SCR-fed worms.**

**(A)** Wild-type animals and *daf-2(e1370)* mutants have increased naïve chemotaxis to 10% butanone with age. Mean ± SEM. n = 10-15 per genotype. ns, not significant (p > 0.05). **(B)** *daf-2(e1370)* mutants maintain learning and **(C)** ITAM abilities better than wild-type animals with age across Days 1, 3 and 5 of adulthood. Mean ± SEM. n = 10-15 per genotype. *p < 0.05, ****p < 0.0001; ns, not significant (p > 0.05).

**(D)** Wild-type animals treated with SCR or INS-17 at either the L4 stage or at **(E)** Day 3 of adulthood had no detectable phenotypic effects in naïve chemotaxis to 10% butanone at Day 5 of adulthood Mean ± SEM. n = 15 per genotype. ns, not significant (p > 0.05).

**(F-G)** Replicates 2 and 3 of data represented in **Fig. 3G.** Probability of survival analysis demonstrates there is no significant difference in lifespan comparing wild-type animals and *ins-17(tm790)* animals. Mean/median ± SEM. n ≥ 100 per replicate. ns, not significant (p > 0.05). **(H-I)** Replicates 2 and 3 of data represented in **Fig. 3H.** Probability of survival analysis for wild-type animals treated with SCR peptide compared to wild-type animals fed INS-17 peptide shows no significant difference in average lifespan. Mean/median ± SEM. n ≥ 100 per replicate. ns, not significant (p > 0.05).

**S4: *daf-16(mu86);ist-1(ok2706)* demonstrate significant memory deficits likely not due to slight sensory deficits in baseline attraction to appetitive butanone.**

**(A)** Wild-type animals and *daf-16(mu86)* mutants have naïve chemotaxis behavior to 10% butanone that is not significantly different. Mean ± SEM. n = 10 per genotype. ns, not significant (p > 0.05). **(B)** SCR and INS-17 peptide treatments have no detectable effects on naïve chemotaxis behavior of wild-type worms nor *daf-16* mutants to 10% butanone. Mean ± SEM. n = 15 per genotype. ns, not significant (p > 0.05). **(C)** Confirmation for experiments in **Fig. 4B** that INS-17 peptide feeding resulted in wild-type animals fed INS-17 to display memory behavior at two hours compared to animals fed SCR, which do not display memory behavior at two hours (forgetting). Mean ± SEM. n = 15 per genotype. **p < 0.01, ****p < 0.0001; ns, not significant (p > 0.05). **(D)** Wild-type animals and *ist-1(ok2706)* mutants have naïve chemotaxis behavior to 10% butanone that is not significantly different. Additionally, **(E)** *ist-1(ok2706)* naïve chemotaxis to attractive 0.1% butanone, while significantly higher than wild-type, is not impaired. Mean ± SEM. n = 15 per genotype. **p < 0.01; ns, not significant (p > 0.05). **(F)** SCR and INS-17 peptide treatments have no detectable effects on naïve chemotaxis behavior of wild-type worms nor *ist-1* mutants to 10% butanone. Mean ± SEM. n = 15 per genotype. ns, not significant (p > 0.05). **(G)** Confirmation for experiments in **Fig. 4D** that INS-17 peptide feeding resulted in wild-type animals fed INS-17 to display memory behavior at two hours compared to animals fed SCR, which do not display memory behavior at two hours (forgetting). Mean ± SEM. n = 15 per genotype. *p < 0.05, ****p < 0.0001; ns, not significant (p > 0.05). **(H)** While *daf-16(mu86); ist-1(ok2706)* double mutants have lowered baseline naïve chemotaxis to the appetitive concentration of butanone (0.1%), these animals do demonstrate attraction to appetitive butanone compared to the neutral concentration (**Fig. S2I**). This suggests that double mutants do not have sensory deficits in attraction to butanone that would prevent them from forming a positive association with the odorant. Mean ± SEM. n = 15 per genotype. ***p < 0.001. **(I)** SCR and INS-17 peptide treatments have no detectable effects on naïve chemotaxis behavior of wild-type worms, *daf-16(mu86)*, *ist-1(ok2706)*, and *daf-16(mu86); ist-1(ok2706)* mutants to 10% butanone. Mean ± SEM. n = 15 per genotype. ns, not significant (p > 0.05). **(J)** Performance indices for STAM and **(K)** forgetting timepoints corresponding to experiments shown in **Fig. 4E-F**. At the T30 timepoint, *ist-1(ok2706)* animals and *daf-16(mu86)* animals fed INS-17 had better memory performance compared to SCR-fed controls, while *daf-16(mu86);ist-1(ok2706)* double mutants displayed no memory behavior. Additionally, wild-type animals and *ist-1(ok2706)* animals fed INS-17 display memory behavior at 2hrs, confirming that peptide feeding worked effectively for these experiments. Mean ± SEM. n = 15 per genotype. ***p < 0.001, ****p < 0.0001; ns, not significant (p > 0.05).

**S5: One-hour food-deprivation does not enhance learning and STAM performance, but does result in increased *ins-17* mRNA transcript levels that may drive more robust ITAM performance.**

**(A)** There are no detectable differences in naïve chemotaxis to 10% butanone in wild-type animals, *ins-17* CRISPR KO animals (*ins-17(knu1320)*), nor *ins-17p::ins-37* (*ins-17(knu1330[ins-37])*) animals. Mean ± SEM. n = 15 per genotype. ns, not significant ($p > 0.05$). **(B)** qRT-PCR of *ins-17* mRNA levels in Day 1 adult wild-type animals show that two hours of food-deprivation results in increased *ins-17* transcript levels. Mean ± SEM. n = 5 per genotype. *$p < 0.05$. **(C)** Wild-type animals FD or non-FD prior to training showed no significant differences between learning and **(D)** STAM behaviors. Mean ± SEM. n = 15 per genotype. ns, not significant ($p > 0.05$). **(E)** Food-deprivation prior to training (1 CS-US pairing) does not enhance *daf-2(e1370)* learning performance, similar to wild-type animal performance. Mean ± SEM. n = 15 per condition. ns, not significant ($p > 0.05$). **(F)** Wild-type animals treated with either SCR or INS-17 peptide had normal naïve chemotaxis prior to either the food-deprivation step or the one-hour feeding period prior to 1 CS-US pairing. These animals also had **(G)** learning and **(H)** STAM behaviors not significantly different between the FD SCR or INS-17 fed groups and the non-FD SCR or INS-17 fed groups. Meanwhile, **(I)** FD wild-type animals that were fed SCR had ITAM performance similar to INS-17 fed counterparts. Mean ± SEM. n = 10-15 per genotype. ns, not significant ($p > 0.05$). **(J)** *ins-17(tm790)* animals have no detectable deficits in chemotaxis to the highly attractive 1% diacetyl compared to wild-type animals. Mean ± SEM. n = 10 per genotype. ns, not significant ($p > 0.05$).

**Table S1.** ***ins-17*** **transcripts increase in response to multiple starvation paradigms and genetic models of nutrient deprivation.**

| Publication Dataset | Experimental Conditions | Result |
|---|---|---|
| Matsunaga et al. 2012 | mRNA expression analysis measured in L2 wild-type, *daf-2(e1370)*, and *daf-16(mu86)* animals | *ins-17* transcripts increased 2.1-fold in *daf-2* animals<br><br>*ins-17* transcripts decreased .34-fold in *daf-16* animals |
| Matsunaga et al. 2012 | *ins-17p*::VENUS expression measured in *daf-2(e1370)* and *daf-16(mu86)* mutants | *ins-17p::*VENUS expression was upregulated in *daf-2* animals and decreased in *daf-16* animals |
| Mueller et al. 2014 | Microarray using RNA from L1 wild-type animals after a 4hr starvation period | *ins-17* transcripts increased after starvation period |
| Chen and Baugh, 2014 | mRNA expression analysis measured during entry into and recovery from L1 arrest | *ins-17* transcripts increased after starvation period<br><br>*ins-17* transcripts decreased after feeding recovery period |
| Hibshman et al. 2017 | Microarray using RNA from L1 wild-type animals 2-4hrs after hatching in the presence or absence of food | *ins-17* transcripts increased in animals that hatched in the absence of food |
| Hibshman et al. 2017 | Microarray using RNA isolated from L1 *daf-16(mgDf50)* animals 2-4hrs after hatching in the presence or absence of food | *ins-17* transcripts increased in *daf-16* animals that hatched in the absence of food |
| St. Ange et al. 2024 | Single nucleus sequencing of neurons in Day 1 adult wild-type and *daf-2* animals | *ins-17* neuronal transcripts are increased in *daf-2* animals compared to wild-type |
| Ogawa et al. 2024 | RNA sequencing of L1 arrested wild-type animals after 4hr period of feeding | *ins-17* transcripts decreased after feeding compared to animals that continued to starve |
